# Microbial eukaryotic predation pressure and biomass at deep-sea hydrothermal vents: Implications for deep-sea carbon cycling

**DOI:** 10.1101/2023.08.11.552852

**Authors:** Sarah K. Hu, Rika E. Anderson, Maria G. Pachiadaki, Virginia P. Edgcomb, Margrethe H. Serres, Sean P. Sylva, Christopher R. German, Jeffrey S. Seewald, Susan Q. Lang, Julie A. Huber

## Abstract

Deep-sea hydrothermal vent geochemistry shapes the foundation of the microbial food web by fueling chemolithoautotrophic microbial activity. Microbial eukaryotes (or protists) play a critical role in hydrothermal vent food webs as consumers, hosts of symbiotic bacteria, and as a nutritional source to higher trophic levels. We measured cell abundances and predation pressures of vent-associated microbial eukaryotes in low temperature diffuse hydrothermal fluids at the Von Damm and Piccard vent fields along the Mid-Cayman Rise in the Western Caribbean Sea. We present findings from experiments performed under *in situ* pressure that show higher cell abundances and grazing rates compared to those done at 1 atmosphere (shipboard ambient pressure); this trend was attributed to the impact of depressurization on cell integrity. A relationship between protistan grazing rate, prey cell abundance, and temperature of end member hydrothermal vent fluid was observed at both vent fields, regardless of experimental approach. The quantification of protistan biomass and grazing pressure shows that hydrothermally-fueled microbial food webs play a significant role in the broader deep-sea carbon budget by contributing to local carbon export and supply of nutrient resources to the deep ocean.

## Introduction

The microbial food web at deep-sea hydrothermal vents is fueled by primary production that is sourced from chemolithoautotrophic microorganisms interacting with diffuse vent fluids. Due to the localized abundance of energy, hydrothermal vent sites support a rich microbial and animal community [1, 2]. Genetic studies have revealed these sites to host highly diverse and distinct bacteria, archaea, viral, and protistan assemblages [3–9]. Unicellular microbial eukaryotes (or protists) are key components of this ecosystem and impact hydrothermal food webs as grazers of local microbial communities [10, 11], parasites [12], hosts to symbiotic bacteria or archaea [13, 14], and a nutritional resource for higher trophic levels (*e.g.,* other protists, mesozooplankton, or invertebrates) [15, 16].

Our understanding of the trophic exchange and flux of nutrients during deep-sea microbial interactions is limited due to the logistical challenges of accurately measuring microbial community interactions *in situ* [17, 18]. The process of collecting vent fluid samples and bringing them shipboard from the deep sea, via CTD casts or vehicle operations, undoubtedly introduces sampling artifacts due to changes in pressure, temperature, and chemical environment [19]. Approaches to reduce sampling bias include instrumentation that enables experimentation at the seafloor or the ability to chemically fix organisms at depth, thereby preserving *in situ* metabolic information [8, 17, 20, 21]. Other methods use chambers that will recover deep-sea fluid and the organisms they contain, while retaining *in situ* pressure during shipboard recovery and subsequent experimental processing [22, 23].

Here, we report measurements of protistan grazing activity and biomass from low-temperature diffuse hydrothermal vent fluids collected from two vent fields that are situated 20 km apart at the Mid-Cayman Rise: the Von Damm and Piccard vent fields. Protistan grazing experiments were conducted at both ambient (shipboard) and *in situ* (using Isobaric Gas Tight chambers - IGTs, [22]) pressure to evaluate how depressurization influences the results of incubations, and to assess the impact that the local hydrothermal vent geochemistry has on microbial biomass and grazing pressure. This study complements previous molecular-based observations of the highly diverse and spatially distinct protistan populations found at the Mid-Cayman Rise [6] with the first assessments of deep-sea hydrothermal vent protistan cell concentration and biomass. Results contribute to efforts in quantifying deep-sea hydrothermal food web interactions, especially those involving microbial eukaryotes.

## Materials and Methods

### Mid-Cayman Rise sample collection

Samples and experiments were collected and executed during cruise AT42-22 (doi: 10.7284/908847) aboard the RV *Atlantis* with ROV *Jason* in January-February 2020 at the Von Damm (2300 m; 18°23’N, 81°48’W) and Piccard (5000 m; 18°33’N, 81°43’W) hydrothermal fields located along the Mid-Cayman Rise. Fluids for shipboard grazing experiments and biogeochemistry were obtained in 10L volume bags (Kynar, Keika Ventures; polyvinylidene fluoride) using the Hydrothermal Organic Geochemistry (HOG) sampler mounted on ROV *Jason* [24]. Between 4-10 L of vent fluid was collected and filtered through a 47 mm polyethersulphone (PES) filter (Millipore) with a pore size of 0.2 µm and preserved with RNAlater (Ambion) at the seafloor for molecular analysis of microbial communities [8]. Fluids for experiments conducted at *in situ* pressure and parallel geochemical measurements were collected with Isobaric Gas Tight (IGT) fluid samplers [22], which filled at a rate of approximately 1 ml sec^-1^, as described previously [6, 10]. Shipboard, dissolved hydrogen gas and methane concentrations were determined by gas chromatography and pH25°C was measured at room temperature with a combination Ag/AgCl reference electrode. Magnesium was measured in a shore-based laboratory by ion-chromatography on stored 30 ml fluid samples.

Non-vent samples were collected from within the overlying non-buoyant hydrothermal plume at each site and from background seawater via CTD-mounted Niskin bottles. Plume samples were identified using *in situ* CTD sensors to detect the presence of hydrothermal influence in real-time (back-scatter and temperature) above each vent field. Background seawater samples were collected outside of the influence of the hydrothermal vent at approximately the same depth as the vent sites (∼2350 m and ∼4950 m; Table S1).

### Microeukaryote grazing experiments

Short-term grazing experiments were conducted using fluids from the Von Damm and Piccard vent fields, from their respective buoyant plumes, and from background seawater collected at depths appropriate for each site (n = 14). The majority of experiments were performed shipboard (at ambient pressure) and a subset were carried out at *in situ* pressures in IGTs for comparison (Table 1). Fluorescently-labeled prey (FLP) uptake experiments were conducted using a modified protocol [25, 26]. For all grazing experiments, FLP, consisting of 5-(4,6-Dichlorotriazinyl) Aminofluorescein (DTAF)-stained and heat-killed *Hydrogenovibrio* [27, 28] was introduced as the analog prey. For additional details on creating FLPs see [10].

**Table 1.** Grazing experiments conducted at the Mid-Cayman rise included 9 from Von Damm and 4 from Piccard. Out of 14 total grazing assays, 5 were conducted in Isobaric Gas Tight chambers at in situ pressure. Temperature reflects the highest recorded temperature at time of fluid collection. Prokaryotic cell concentrations were derived from discrete fixed samples from the same fluid, while eukaryotic cell concentrations reported below are derived from the T0 grazing experiment time points. When prokaryote cell count samples were not countable (due to mineral precipitation in the sample), an average prokaryotic cell ml^-1^ was used for downstream calculations (7.11 x 10^4^ cell ml^-1^). Absent grazing rates include those that had a negative slope and percentage prokaryote turnover shows the relative top-down (higher percentage) to bottom-up pressures on the microbial communities, based on grazing rate and cell concentrations.

| Vent field | Site name | Fluid origin | Fluid ID | Average eukaryote cells $\text{ml}^{-1}$ | Eukaryote cells $\text{ml}^{-1}$ (min / max) | Average prokaryote cells $\text{ml}^{-1}$ | Prokaryote cell $\text{ml}^{-1}$ (min / max) | Rate $\text{min}^{-1}$ | Cells consumed $\text{ml}^{-1} \text{ hr}^{-1}$ | mL grazer$^{-1} \text{ hr}^{-1}$ | % removed prokaryotes $\text{day}^{-1}$ | Temperature ($^{\circ}\text{C}$) |
| --- | --- | --- | --- | --- | --- | --- | --- | --- | --- | --- | --- | --- |
| Von Damm | Background | CTD002 | Niskin 8,10 | 91.838 | 69.97 / 113.7 | $3.79 \times 10^4$ | $1.8 / 5.6 \times 10^4$ | 0.00296 | 127.07 | $3.7 \times 10^{-5}$ | 8.0 | 4.2 |
| | Plume | CTD001 | Niskin 2 | 157.77 | 55.98 / 284.4 | $1.65 \times 10^4$ | $1.4 / 1.9 \times 10^4$ | 0.00527 | 24.03 | $9.2 \times 10^{-6}$ | 3.5 | 4.2 |
| | Mustard Stand | J2-1243 | LV17 | 259.77 | 230.9 / 288.6 | $5.67 \times 10^4$ | $4.2 / 7.1 \times 10^4$ | bd | bd | bd | bd | 108.0 |
| | Old Man Tree | J2-1238 | IGT4 | 349.86 | -- | -- | -- | 0.00376 | 1050.4 | $4.2 \times 10^{-5}$ | 35.0 | 121.6 |
| | Ravelin #2 | J2-1238 | LV13a | 409.33 | 335.9 / 482.8 | -- | -- | 0.00347 | 116.86 | $4.0 \times 10^{-6}$ | 3.9 | 94.0 |
| | Ravelin #2 | J2-1244 | IGT4 | 944.62 | -- | -- | -- | 0.02106 | 15867.1 | $2.4 \times 10^{-4}$ | 540.0 | 98.2 |
|  | Ravelin #2 | J2-1244 | IGT5 | 629.74 | -- | -- | -- | bd | bd | bd | bd | 98.2 |
| | Shrimp Hole | J2-1244 | LV13 | 385.72 | 377.8 / 393.6 | $4.2 \times 10^4$ | $3.9 / 4.5 \times 10^4$ | bd | bd | bd | bd | 21.0 |
| | X-18 | J2-1235 | LV23, Bio5 | 314.87 | 209.9 / 419.8 | $1.11 \times 10^5$ | $1.1 / 1.1 \times 10^5$ | 0.00174 | 1166.3 | $3.3 \times 10^{-6}$ | 25.0 | 48.0 |
| Piccard | Plume | CTD004 | Niskin 10 | 79.301 | 55.98 / 112 | $5.14 \times 10^4$ | $4.7 / 5.6 \times 10^4$ | 0.00536 | 44.26 | $1.1 \times 10^{-5}$ | 2.1 | 4.5 |
| | Lots 'O Shrimp | J2-1241 | LV24 | 230.91 | 230.9 / 230.9 | $5.39 \times 10^4$ | $2.6 / 9.0 \times 10^4$ | bd | bd | bd | bd | 36.0 |
| | Shrimpocalypse | J2-1240 | LV13 | 454.82 | -- | $2.39 \times 10^5$ | $1.1 / 3.2 \times 10^5$ | 0.01569 | 6006.91 | $5.5 \times 10^{-5}$ | 60.0 | 85.0 |
| | Shrimpocalypse | J2-1240 | IGT3 | 384.84 | -- | $2.39 \times 10^5$ | $1.1 / 3.2 \times 10^5$ | 0.01679 | 17284.73 | $1.9 \times 10^{-4}$ | 170.0 | 85.0 |
| | Shrimpocalypse | J2-1240 | IGT7 | 524.79 | -- | $2.39 \times 10^5$ | $1.1 / 3.2 \times 10^5$ | n/a | n/a | n/a | n/a | 85.0 |

bd indicates not detected or below detection limit for grazing rate
n/a indicates grazing experiment was not 'countable'
-- indicates no value is available
\*For prokaryote cell count samples that were not countable, an average across all vents was taken and used for calculations

#### Incubations conducted at ambient pressure

Large volume bags filled with vent fluid using the HOG sampler were subsampled into 2-3 acid-rinsed and clean bags (polyvinylidene fluoride) at volumes ranging from 1.5-2 L (volume and experimental replicates varied based on available water budget). Each shipboard grazing experiment was conducted in duplicate or triplicate (where all treatment volumes were the same; Table S2) and kept at ∼22°C for the incubations. For control treatments, fluid was filtered through a 0.8 µm porosity filter in duplicate (0.5-1L) to remove microbial predators. Immediately after distributing the experimental and control fluid, thoroughly mixed FLP was introduced to each treatment so the concentration of FLP was 20-25% of the *in situ* prokaryotic community, gently mixed, and a T0 sample was. At each time point, 200 ml of fluid was preserved using chilled formaldehyde (1% final concentration) in darkened amber bottles (20 ml for controls) and stored in the dark at 4°C until processing. Target time points ranged from 0 minutes to 40 minutes (T0, T10, T15, T20, and T40 or Tf); in some cases, T20 time points were not taken due to constraints on recovered hydrothermal fluid volume (Table S2). Following the final time point (Tf), all fluid from experimental bags was filtered into a Sterivex filter with porosity 0.2 µm (Millipore), preserved with RNAlater, and frozen at −80°C for genetic analysis.

#### Incubations performed at in situ pressure

Before deploying each IGT sampler, the dead volume was filled with 0.2 µm-filtered background deep seawater and a Teflon O-ring was added to the sample chamber to enhance mixing of collected fluid and injected amendments. ROV Jason positioned the IGT inlet with a co-located temperature probe to collect diffuse vent fluid. Shortly after ROV recovery, a titanium piston separator was affixed to the IGT [similar to 23] to facilitate the introduction of FLP and collect subsamples for grazing experiment time points without rupturing cells by eliminating the need for fluids to pass through the small opening of a pressure retaining sample valve.

Each IGT-based experiment was maintained at *in situ* pressure (Table 1) for the duration of the incubation using a high performance liquid chromatography (HPLC) pump to compensate for pressure loss during subsampling [also see 23]. FLPs were premixed at a final volume of 8 ml to add to the 150 ml volume of the IGT samples and the final FLP concentration was 1 x 10^4^ cells ml^-1^. After agitating the IGT chamber to gently mix the collected vent fluid samples with the added FLPs, an initial (T0) time point was taken by moving the sample into the pressure separator and then emptying a 30 ml sample into amber bottles with chilled formaldehyde (final concentration 1%). Time points were planned for 0, 10, 20, and 40 minutes, but time constraints meant that time points were often taken at irregular intervals (compared to the shipboard incubation sample intervals; Table S2). Control treatments concurrent with the IGT samples from vents were not feasible, IGT control treatments were conducted separately by filling IGTs with 0.8 µm filtered deep-sea background seawater (collected via CTD), bringing the IGT chamber to *in situ* pressure (3000-6000 psi), and repeating the FLP experiment procedure.

Since opportunities for biological replicate incubations were limited (only 2; Table S2), technical replicate cell counts were completed to provide additional confidence in our findings (repeat microscopy counts). Results from technical replicates are reported in the Supplementary Information.

### Processing grazing experiment samples

Formaldehyde-fixed samples (final concentration 1%) were kept in the dark and at 4°C until analysis for both prokaryotic and eukaryotic counts. For the *in situ* prokaryotic cell concentration (bacteria and archaea, or microbial prey population), between 1-10 ml was filtered onto 0.2 µm black polycarbonate filters and counted under epifluorescence (blue/cyan filter for DAPI-stained cells). Similarly, 2-5 ml of the grazing experiment control samples were filtered onto 0.2 µm black polycarbonate filters and counted under the fluorescein isothiocyanate (FITC) filter to ensure the number of FLP did not change for the duration of the experiment. Samples for all grazing treatments were filtered onto 0.8 µm black filters (volumes ranged between 100-200 ml) and stained with a DAPI solution (final concentration of DAPI ∼10 μg/ml). Filters for DAPI and FITC were used to count the number of nano-(< 20 µm) and micro-(≥ 20 µm) eukaryotic cells observed and the number of FLP inside each eukaryotic cell (by switching back and forth). This approach enabled the enumeration of the total number of eukaryotic cells ml^-1^ and the number of ingested FLP (Table S3). A minimum of 30 fields of view were counted for each sample at 100x magnification. Eukaryotic cells were distinguished from other DAPI-stained debris by noting the presence of a nucleus or eukaryote-like cell morphologies (*e.g.*, flagella, cilia, or organelles).

#### Quantifying protistan predation and biomass

Microscopy counts revealed the number of FLPs ingested per eukaryotic cell, concentration of bacteria and archaea, and concentration of microbial eukaryotes. For each grazing assay, the slope of the best fit line of the average number of ingested FLPs by eukaryotes vs. incubation time was calculated. At each time point, the mean number of FLPs ingested per total eukaryotes observed and the standard mean error across replicate experiments was determined. The slope of the best fit line was estimated. Due to the small volume of the IGT experiments and the observation that the final IGT time point (Tf) varied drastically from other time points, the IGT Tf samples were removed before estimating the slope. The slope of the best fit line equates to the number of FLPs consumed by a protistan grazer every minute. Following established protocols [29], the clearance rate (ml grazer^-1^ hr^-1^) and grazing rate (cells consumed ml^-1^ hr^-1^) were calculated (Table S3). Grazing experiments that resulted in negative slopes were interpreted as ‘undetected’ or ‘below the detection limit’ grazing; these experiments are presented as 0 in results. See Supplementary Table S3 for a list of calculations and related references for quantifying grazing impact.

Short-term FLP uptake experiments also provided the visualization of eukaryotic cells. Zeiss image processing software determined the “height” and “width” of preserved cells where height equates to the longest dimension and width equals the longest cross section [30]. Eukaryotic cells used for biovolume calculations were imaged from a random assortment of experimental time points; the volume of each cell was determined (µm^3^) based on equations from Pernice et al. [31] and Hillebrand et al. [32]. Biovolumes were converted to carbon cell^-1^ using carbon conversion rates from Meden-Deuer and Lessard [33] (Table S3). In these studies the carbon content of several heterotrophic protists ranged from 4.7 x 10^3^ – 1.2 x 10^7^ µm^3^. We considered estimates from Meden-Deuer and Lessard [33] to represent the overall range of likely carbon content for heterotrophic cells, assuming that protistan cells captured in our samples are largely heterotrophic. Further, we acknowledge that these estimates are based on heterotrophic species in culture, which are likely physiologically distinct from cells originating from deep-sea vents. Secondly, we calculated carbon content per cell by incorporating field standard estimates for nano-and micro-eukaryotes ([34], see Table S3). Since this work is derived from preserved cells that may have shrunk [35, 36] or have gone through depressurization, both measured biovolume and a range of pg C cell^-1^ was considered to present a range of carbon conversion approaches (see Table S3). Similarly, we included a range of carbon conversion factors for prokaryotes [37, 38]. The range of carbon content cell^-1^ was used to constrain microbial eukaryotic and prokaryotic pools in all calculations.

### Amplicon sequence survey

Filters retrieved from ROV *Jason* and from the final time point of each grazing assay were processed identically. RNA was extracted from frozen filters (stored in RNAlater) as amplicon sequences originating from extracted RNA are more likely to represent metabolically active cells, rather than inactive cellular material that may have sunk from above. The filter was first separated from the RNAlater and distributed into tubes with a lysis buffer (Qiagen 1053393). The RNAlater was centrifuged for 15 minutes at 16,000 x g, and the supernatant was removed. Lysis buffer was added on top of any cellular materials collected, vortexed, and then combined with the filter. The filter and lysis buffer solution was vortexed thoroughly with RNAase-free silica beads. The lysis buffer was then separated from beads and filter material with a syringe and processed using the Qiagen RNeasy extract kit (Qiagen 74104), which included an in-line RNAse-free DNase removal step (Qiagen 79256). Total RNA was reverse transcribed to cDNA and amplified with V4-specific primers [39]. MiSeq 2 x 300 bp PE sequencing was performed at the Marine Biological Laboratory Bay Paul Centre Keck sequencing facility.

Amplicon sequences were processed using QIIME2 (version 2021.4; [40] as described in [6]. Following quality control, Amplicon Sequence Variants (or ASVs) were determined from the sequences and taxonomic assignment was done using the PR2 database (v 4.14; [41, 42]. For this analysis, we focused primarily on the microeukaryotic population, removing sequences assigned to prokaryotes or Metazoa. Similar to [6], ASVs were categorized as vent-only or cosmopolitan, based on their presence in only vent samples or throughout vent, plume, and background samples, respectively.

*In situ* samples and Tf samples were compared to subset for taxa that may have been enriched within each grazing experiment. If an ASV was present in both *in situ* and associated Tf samples, it was considered a member of the captured protistan community. If the total number of sequences and or ASVs within the group increased, then the taxonomic group was considered to be enriched. In another approach to determine which taxa may be enriched across the vent sample types, we employed the corncob analysis [43]. Corncob models the relative abundances and the differential abundances of the ASVs as a linear function of vent habitat versus non-vent habitat. Using the parametric ‘Wald Test’, corncob allows us to test the hypothesis that a given ASV will change significantly across the parameters. Positive coefficients indicate that, at the Family level, the taxonomic group was enriched in vent samples compared to the non-vent samples.

## Data availability

Intermediate data products and required code to reproduce results can be found at https://shu251.github.io/midcayman-rise-microeuk/. Raw sequence data are available through NCBI SRA BioProject accession number PRJNA802868.

## Results

### Fluid geochemistry

The Von Damm and Piccard hydrothermal vent fields at the Mid-Cayman Rise are located at different depths, 2350 m and 4950 m, respectively, where Piccard is the deepest known hydrothermal vent field [44, 45]. At the time of sample collection, low temperature diffuse fluid from Von Damm ranged between 12-129°C, while temperatures at Piccard were 19-85°C (Table 1; Table S1). Fluid venting from Von Damm is largely influenced by ultramafic rock, and contains less dissolved sulfide, and has higher concentrations of methane compared to fluids from Piccard. Mafic rock at Piccard results in vent fluids that are more acidic and enriched in dissolved sulfide; Piccard is enriched in dissolved hydrogen, compared to other hydrothermal systems, as a result of the high pressure, temperature, and basalt-hosted system [46, 47] (Table S1).

### Microbial biomass

Cell abundances for both bacteria and archaea (prokaryotes) and eukaryote populations shared a similar trend where the highest concentration (cell ml^-1^) was found within diffuse vent fluids, followed by the plume and background environments (Figures 1a-b). Non-vent (plume and background) prokaryote cell concentrations averaged 3.5 x 10^4^ cells ml^-1^, while concentrations within diffuse vent fluids averaged 1.4 x 10^5^ cells ml^-1^. Prokaryotic cell concentrations within Piccard diffuse fluid (1.9 x 10^5^ cell ml^-1^) were higher than those at Von Damm (7.0 x 10^4^ cells ml^-1^; Table 1; Figure 1b).

**Figure 1.**
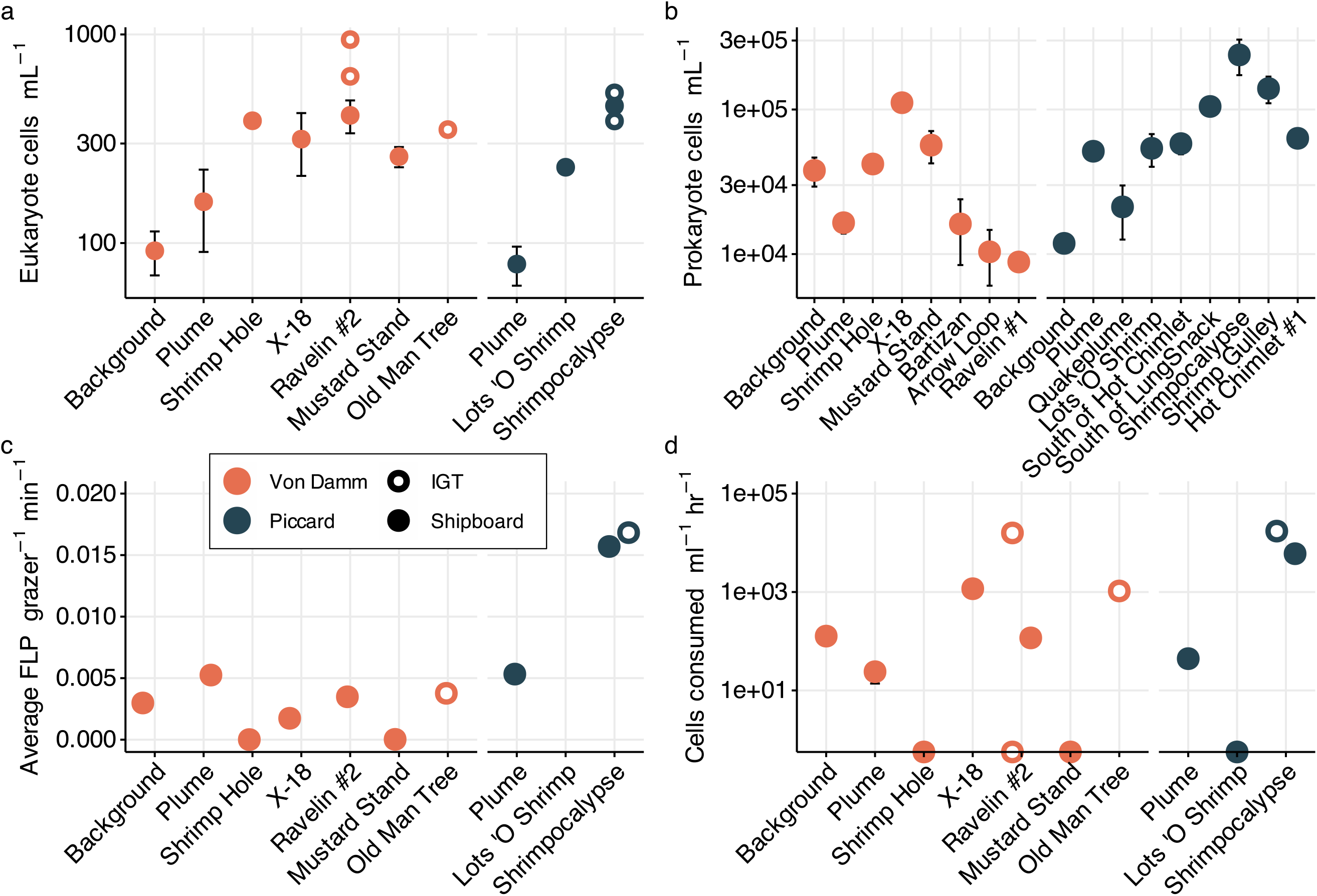
Cell concentration and grazing results from Mid-Cayman Rise. (a) Eukaryote cells ml^-1^ (log scale) at time zero for each experiment. Error bars represent standard mean error. (b) Cell concentration of bacteria and archaea (cells ml^-1^; log scale). (c) Average number of FLPs observed inside a eukaryotic cell per minute, derived from grazing experiment cell counts. And (d) grazing rate, expressed as the number of cells consumed by protistan predators ml^-1^ hr^-1^ (log scale). Symbol color denotes vent field, filled in circles are derived from shipboard experiments or samples (ambient pressure), while circle outlines represent results from IGT experiments (*in situ* pressure). Vent sites along the x-axis are arranged from lowest to highest temperature (left to right) for Von Damm and Piccard)

Eukaryote cell concentrations within the background and plume environments averaged 1.1 x 10^2^ cells ml^-1^. Within diffuse vent fluids, eukaryotic cell abundances were higher than non-vent environments, averaging 3.7 x 10^2^ cells ml^-1^. Eukaryotic cell concentrations were similar between Piccard and Von Damm, averaging 4.0 x 10^2^ cells ml^-1^ and 3.2 x 10^2^ cells ml^-1^, respectively. (Table 1; Figures 1a-b). The average eukaryotic cell concentrations derived from samples collected with IGTs, and thus maintained at *in* situ pressure, were slightly higher than samples collected with the HOG sampler and then used for shipboard experiments, 4.5 x 10^2^ cells ml^-1^ versus 3.3 x 10^2^ cells ml^-1^. Values reported here include the total number of protists counted (both nano-and micro-size classes captured on 0.8 µm filters); results from eukaryotic cell counts separating nano-, micro-, and total (nano + micro) are reported in Figure S1. Since the process of fixation can shrink cell volume [35, 36] and depressurization may have impacted cell integrity, our distinctions of micro-versus nano-plankton size classes may not be accurate. Additional supporting evidence for this can be found in the Supplemental Information. The majority of our downstream results consider the total microeukaryote population.

We incorporated both biovolume-specific and field standard estimates of carbon biomass for the microbial eukaryotic population. This enabled us to factor in potential downstream impacts of preservation and depressurization on cell morphology. The average biovolume for protists counted outside of vent fluid was 773 µm^3^ and 3321 µm^3^ for protistan cells found within vent fluids (Table S4; [32]). Biovolume derived from shipboard results averaged only 1976 µm^3^, compared to over 4000 µm^3^ from the IGT results (average across IGT and shipboard results was used for estimated C cell^-1^). Using a field standard carbon conversion rate of 0.216 pg C cell^-1^ volume^0.939^ [33], we determined a putative pg C cell^-1^ value for the vent and non-vent associated cell abundances. The non-vent background seawater microeukaryote population averaged 109.2 pg C cell^-1^, while cell abundances within diffuse vent fluids averaged 414.8 pg C ml^-1^ (Table S5). This equates to an estimated total carbon pool of 12.9 µg C L^-1^ outside of diffuse vent fluids and 172 µg C L^-1^ within hydrothermal vent fluids (Table 2). As an alternative to biovolume, we used carbon conversion factors specific for micro-and nanoplankton, 138 pg C cell^-1^ and 2.6 pg C cell^-1^, respectively [34]; also see Tables S3 and S5). Similar to the biovolume-derived estimation for carbon, the estimated carbon pool of microeukaryotes within diffuse vent fluids was higher, 5.0 µg C L^-1^, compared to the non-vent microeukaryote population, 2.5 µg C L^-1^ (Table 2). Similarly, both biovolume-derived and size-fraction derived carbon biomass estimates were higher among samples from IGT experiments compared to samples from experiments at ambient pressure (Table 2). The range of carbon biomass estimates by experiment are also reported in Figure S2.

**Table 2.** Summary of microeukaryotic carbon biomass estimated by biovolume or size fraction. Values reported below are the mean across background and plume (non-vent) or diffuse vent fluid only, or between vent sites at Piccard and Von Damm.

| **Category** | **Carbon**<br>**conversion**<br>**strategy** | **µg C per L** | **Maximum µg**<br>**C per L** | **Minimum µg**<br>**C per L** |
| --- | --- | --- | --- | --- |
| Non-vent | Biovolume | 12.9 | 17.2 | 8.7 |
| Vent |  | 172.2 | 391.8 | 38.1 |
| Piccard (vent only) |  | 165.4 | 217.7 | 95.8 |
| Von Damm (vent only) |  | 175.6 | 391.8 | 38.1 |
| Shipboard (vent only) |  | 127.2 | 188.7 | 38.1 |
| IGT (vent only) |  | 224.9 | 391.8 | 145.1 |
| Non-vent | Size fraction | 2.5 | 3.5 | 1.5 |
| Vent |  | 5.0 | 11.1 | 0.2 |
| Piccard (vent only) |  | 4.6 | 10.8 | 0.6 |
| Von Damm (vent only) |  | 5.2 | 11.1 | 0.2 |
| Shipboard (vent only) |  | 3.5 | 7.8 | 0.2 |
| IGT (vent only) |  | 8.5 | 20.5 | 1.0 |

### Protistan grazing

The observed number of FLPs consumed by protistan grazers throughout each experiment was fit with a best fit line, where the slope represents the average number of FLPs consumed by protistan grazers per minute ([29]; Tables S2-S3; Figure S3). When the slope of the line was negative, grazing was considered undetected or below detection and replaced with a zero value (Figures 1c-d). Since eukaryotic cell abundance decreased over time in these experiments (Figure S4), zero values were not included in reported averages, but are included in Figures 1-2. Eukaryotic cells ml^-1^ dramatically decreased in the time point for each IGT experiment (Tf), warranting the removal of this time point, due to bottle effects (Figure S4). Results from control experiments demonstrated stable FLP concentrations over time (Figure S5).

**Figure 2.**
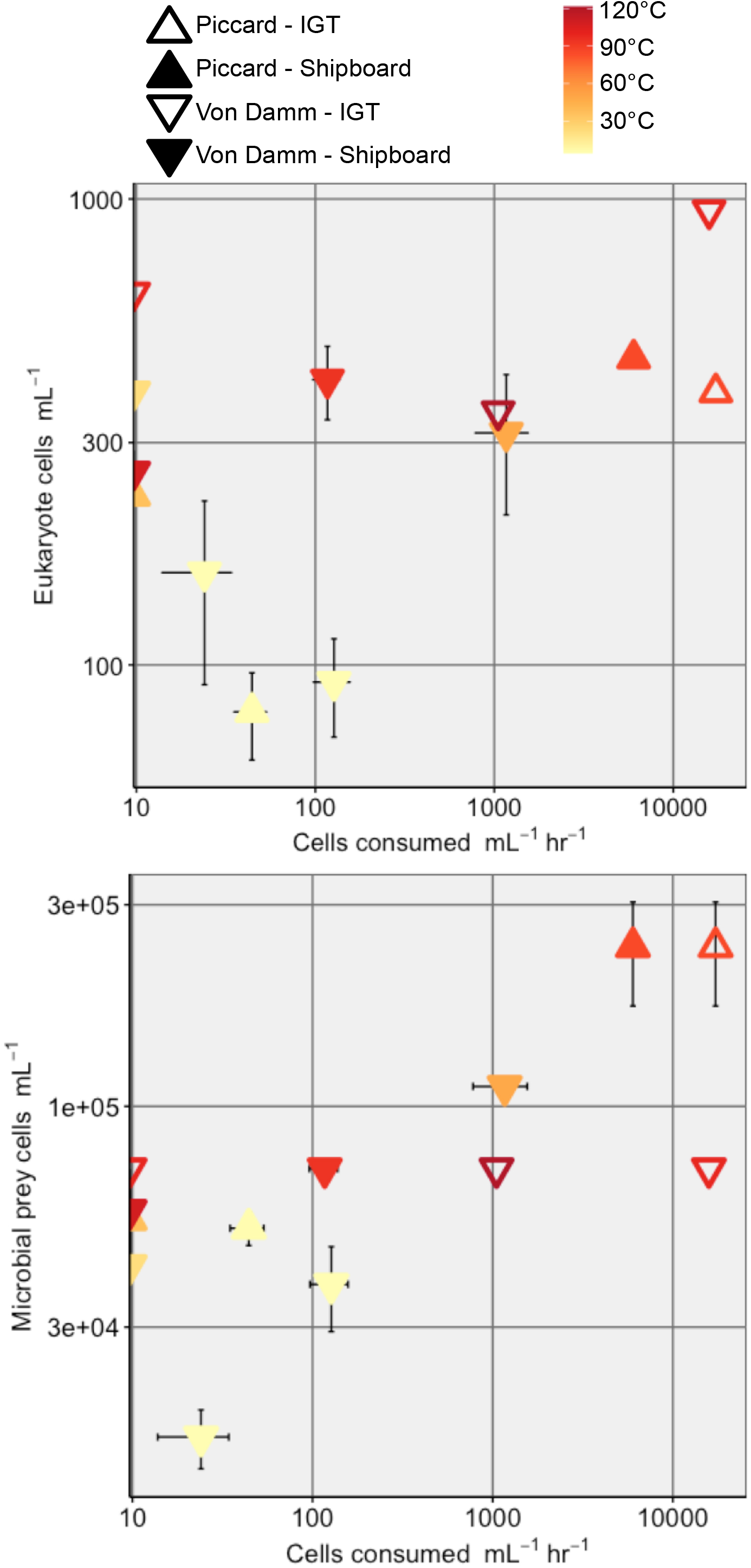
Grazing rate (Cells consumed ml^-1^ hr^-1^) along the x-axis with (a) eukaryote cells ml^-1^ and (b) prokaryote cells ml^-1^ along the y-axis. Symbol color denotes the temperature of fluid at time of sample collection (°C). Filled in triangles are derived from shipboard experiments or samples, while circle outlines represent results from IGT experiments.

The average grazing rate for experiments conducted with diffuse vent fluid was 6.9 x 10^3^ cells consumed ml^-1^ hr^-1^ (min: 116.9, max: 1.7 x 10^4^ cells consumed ml^-1^ hr^-1^), which was higher than non-vent samples, where grazing rate averaged 65 cells consumed ml^-1^ hr^-1^ (min: 24, max: 127 cells consumed ml^-1^ hr^-1^; Figure 1). Between the two vent fields, grazing rates within Piccard diffuse fluids (1.2 x 10^4^ cells consumed ml^-1^ hr^-1^) were higher than Von Damm (4.6 x 10^3^ cells consumed ml^-1^ hr^-1^). While fewer IGT experiments were conducted, the IGT results yielded higher grazing estimates, averaging 1.1 x 10^4^ cells consumed ml^-1^ hr^-1^ at Piccard compared to 2.4 x 10^3^ cells consumed ml^-1^ hr^-1^ at Von Damm. By incorporating biomass estimates of microbial prey (86 fg C cell^-1^; Morono et al. 2011; Table S3), we determined the amount of carbon that may be taken up by the grazer community outside of the vent environment as 5.6 pg C ml^-1^ hr^-1^; based on experiments run at ambient and *in situ* pressure, the rate of carbon consumption within the vent is 209 pg C ml^-1^ hr^-1^ and 980 pg C ml^-1^ hr^-1^, respectively (Table 3).

**Table 3.**
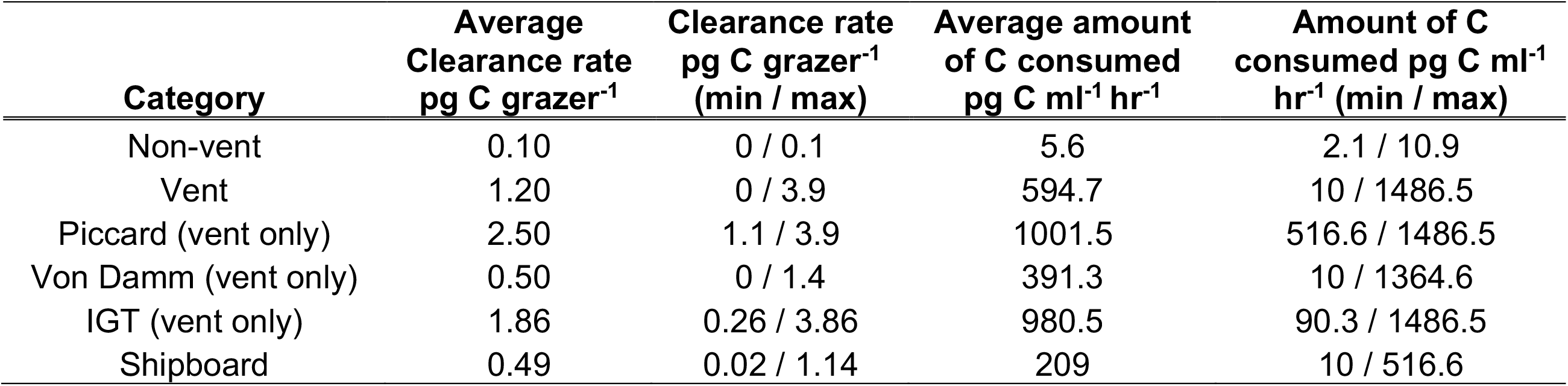
Estimate of the amount of carbon consumed by the protistan grazer population. Using either the clearance rate or grazing rate.

| **Category** | **Average Clearance rate**<br>**pg C grazer<sup>-1</sup>** | **Clearance rate**<br>**pg C grazer<sup>-1</sup>**<br>**(min / max)** | **Average amount**<br>**of C consumed**<br>**pg C ml<sup>-1</sup> hr<sup>-1</sup>** | **Amount of C**<br>**consumed pg C ml<sup>-1</sup>**<br>**hr<sup>-1</sup> (min / max)** |
| --- | --- | --- | --- | --- |
| Non-vent | 0.10 | 0 / 0.1 | 5.6 | 2.1 / 10.9 |
| Vent | 1.20 | 0 / 3.9 | 594.7 | 10 / 1486.5 |
| Piccard (vent only) | 2.50 | 1.1 / 3.9 | 1001.5 | 516.6 / 1486.5 |
| Von Damm (vent only) | 0.50 | 0 / 1.4 | 391.3 | 10 / 1364.6 |
| IGT (vent only) | 1.86 | 0.26 / 3.86 | 980.5 | 90.3 / 1486.5 |
| Shipboard | 0.49 | 0.02 / 1.14 | 209 | 10 / 516.6 |

Grazing rates corresponded to the microbial cell abundances and temperature of fluid sampled at both Von Damm and Piccard (Figure 2). The highest grazing rates (>1000 cells ml ^-1^ hr^-1^) were generally found at sites with higher concentrations of microeukaryotes (> 300 cells ml ^-1^) and microbial prey cells (>1.0 x 10^5^ cells ml^-1^). Additionally, vent field and fluid temperature appeared to play a role in the trend between microbial prey concentration and protistan grazing rate (Figure 2b). The highest protistan grazing rates at Piccard corresponded to the highest concentration of microbial prey and temperature maxima. By contrast, increasing temperatures at Von Damm (beyond 100°C) appeared to limit microbial prey concentration and subsequent grazing rate (Figure 2b). Patterns observed between eukaryotic cell abundance, microbial prey abundance, temperature, and grazing rate were consistent, regardless of the pressure conditions of the incubation (Figure 2). For three experiments in which grazing was deemed undetectable (negative slope), the temperature, vent fluid, and cells ml^-1^ did not show a predictable pattern. Instead, these were attributed to the highly mixed, wafty, and ephemeral nature of the diffuse flow and seawater interface. Comparisons of grazing rate with other environmental parameters were not found to have a relationship (Figure S6).

### Links to species composition

To investigate specific protistan taxonomic groups that may be linked to elevated grazing activity or hydrothermal vent habitat type, as well as how communities changed during the grazing experiments, we compared the community composition, derived from 18S rRNA gene tag-sequence analysis, across vent and non-vent samples and between the *in situ* and Tf samples. Generally, the alveolate taxa, ciliates and dinoflagellates, outnumbered other recovered taxa in both species richness (ASV richness) and sequence number (comparative relative abundance). Second to the alveolates, hacrobia, rhizaria, and members of the stramenopile groups were consistently present across hydrothermal vents at Mid-Cayman Rise (also see [6]). Since relative abundance of 18S rRNA gene amplicons is not representative of cell biomass and gene copies can vary significantly by species, we drew the majority of our observations from transformed data to minimize these artifacts [48, 49].

Ordination analysis revealed that the community composition of protistan communities from diffuse fluid generally clustered with corresponding Tf grazing experiment samples (open versus shaded symbols in Figure 3a). ASVs that appeared in both *in situ* samples and samples from grazing incubations were assumed to represent taxa contributing to grazing; of these ASVs, over 1,500 were found to be shared at Piccard and Von Damm sites and the majority were also cosmopolitan (found at vent and non-vent sites). Comparisons between *in situ* and grazing Tf samples at the ASV level revealed a higher occurrence of dinoflagellates, radiolaria, and opalozoa in non-vent experiments compared to vent sites. Overall, ciliates and dinoflagellates appeared to be the predominant protistan grazers in all experiments.

**Figure 3.**
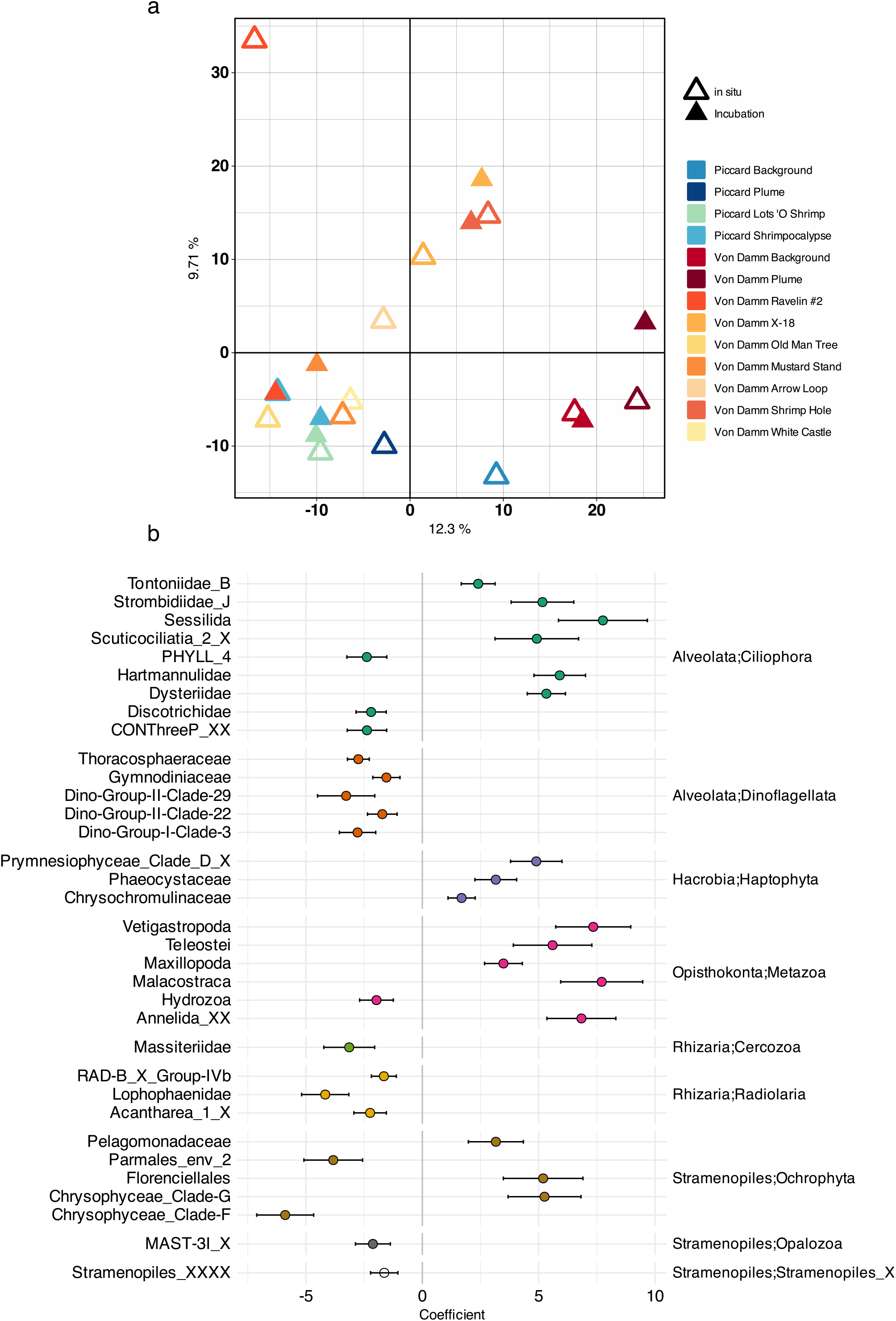
(a) Ordination analysis across all sites sampled and associated Tf for grazing incubations. Color designates each vent site, plume, or background sample and symbol differentiates the vent field. (b) Output from corncob analysis [43] that identified specific families that may be enriched within vent samples (positive coefficient) compared to non-vent samples (negative coefficient; include background and plume).

Positive coefficients derived from the corncob analyses at the family level demonstrated that several protistan families were enriched at vent sites compared to non-vent samples (Figure 3b). These results show that for major taxonomic groups, specific families are enriched within the vent samples; including families within the ciliates, haptophyta, and ochrophyta. For instance, many of the ciliate groups had positive coefficients, such as the *strombidiae*, *scuticociliates*, while most other families did not. Although most dinoflagellate families were not enriched at the vent sites compared to the non-vent samples, their prominence still suggested that they were a key player in the vent protistan community (Figure 3b). Within the stramenopiles, Pelagomonadales, *Dictyochophyceae* and Clade G of Chrysophyceae were the only families to show consistent enrichment at the vent sites.

## Discussion

We quantified microbial eukaryotic cell concentration and predation pressure across two deep-sea hydrothermal vent fields using grazing experiments conducted at both ambient (1 atmosphere) and *in situ* deep-sea pressures. Our study at the Mid-Cayman Rise offers the opportunity to compare two vent fields, Von Damm and Piccard, that are located close together, but at separate depths and have distinct geochemistry. Regardless of vent field, grazing experiments retained at *in situ* deep-sea pressure revealed consistently higher concentrations of eukaryotic cell abundances and predation rates, demonstrating the value in maintaining *in situ* conditions for incubations. We also place our results in the larger context of deep-sea hydrothermal vent food webs, and show a food web with constrained microeukaryotic carbon biomass and exchange (Figure 4).

**Figure 4.**
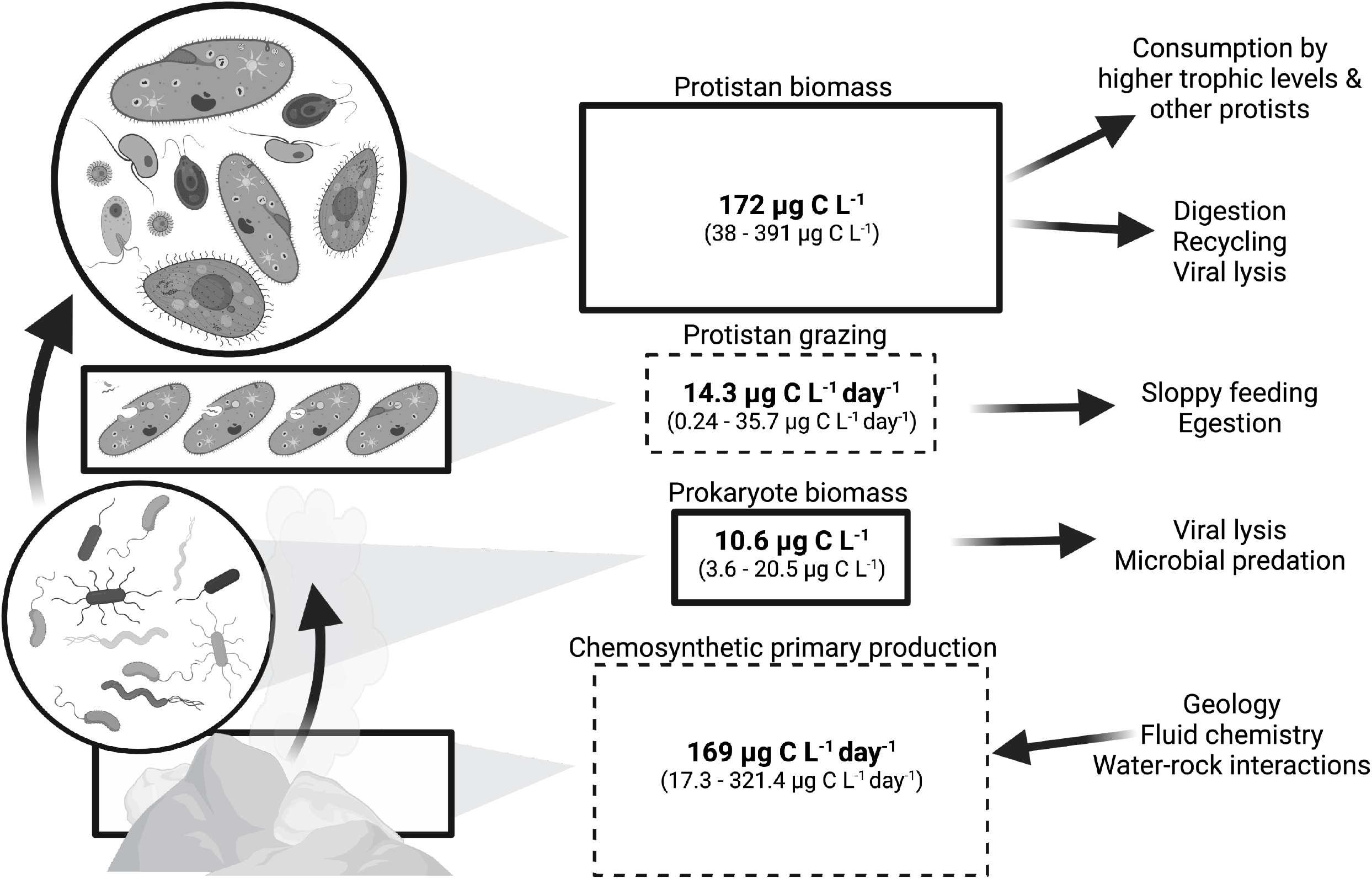
Schematic of the microbial food web at the Mid-Cayman Rise hydrothermal vent fields in terms of carbon. Rates are expressed as µg C L^-1^ day^-1^ (dashed line boxes) and biomass is represented by µg of C L^-1^ (solid line boxes). Arrows show the net flow of carbon to higher trophic levels and unconstrained losses. Created with BioRender.com.

### The importance of deep-sea microbial interactions in situ

Differences between experiments conducted with vent fluid kept in IGTs vs. collected with ROV *Jason* reflect the influence that depressurization likely has on deep-sea protistan survival and activity. Experiments conducted at 1 atmosphere underestimated grazing rates and cell abundances. Biological replicates from Ravelin #2 (Von Damm) and Shrimpocalypse (Piccard) allowed direct comparison of microeukaryote cell abundances between IGT and ambient experiments (Table 1), despite the difference in incubation volume. Eukaryotic cell abundances and carbon biomass at Ravelin #2 and Shrimpocalypse were consistently higher within IGT experiments, which we interpret as *in situ* pressure maintaining cell structure and integrity [17, 19]. Consistent with this observation, average cell abundances (Figure 1a) and biomass (Table 2) for Von Damm were slightly higher compared to Piccard. Since Piccard vent fluid would experience a larger change in pressure during collection, we speculate that this contributed to differences downstream results.

Molecular analyses revealed the microbial eukaryotic community composition to be similar between the *in situ* vent fluid and final time point of each grazing experiment (Figure 3a). This provides evidence that our shipboard experiments largely captured and retained microbial communities representative of the deep sea. Since the majority of ASVs shared between the *in situ* diffuse fluid samples and grazing incubations were found at both vent fields and present throughout the vent and non-vent habitats, we hypothesize that collection and depressurization ahead of the shipboard incubations selected for protists that are more ubiquitous throughout the deep sea (cosmopolitan), rather than isolated to hydrothermal vent sites. Further, many of the selected protists may include barotolerant taxa, a trait that exists in many species, but is highly variable and species-specific [19, 50]. The microbial prey population in our experiments was assumed to be representative of the diffuse vent community. This assumption is derived from previous evidence, where the prokaryotic community remains compositionally similar to those *in situ*, when used in shipboard experiments, while gene expression results show evidence that cells experience environmental stress [21].

Grazing experiments conducted at both pressure conditions show the important relationship between the chemosynthetic and protistan trophic levels, highlighting a critical route of carbon flux within hydrothermal vent food webs. Despite differences in vent fluid origin and pressure condition of the grazing experiments, the same relationship between microbial cell abundance, diffuse vent fluid temperature, and protistan grazing rates was found (Figure 2). Adding to the value of pursuing experiments conducted at ambient and *in situ* pressures, when results from this study are compared with a previous deep-sea vent protistan grazing study that used a different experimental approach [10], grazing rates and minimum-maximum values are comparable (Figure S7).

### Trends in microeukaryotic cell biomass and grazing activity

Protistan top-down pressure varied at separate vent fields and within the same vent field (Figure 1). Individual vent sites (1-10s meters apart) are known to host highly diverse and distinct microbial communities between vent sites [6], which likely contributes to the observed range in grazing rates [51]. Similarly, other studies that measure protistan grazing and biomass often observe a range of values. In particular, a study that used a sampling device to conduct experiments *in situ* reported grazing rates ranging from 18.7-13,600 cells ml^-1^ hr^-1^ [52], which was comparable to our results 24-17,200 cells ml^-1^ hr^-1^. In a coastal ecosystem, Connell et al. [53] reported a range of 1.74 to 28.8 µg C L^-1^ day^-1^ consumed by heterotrophic protists, which is in line with ranges of µg of C consumed in our findings across the Mid-Cayman Rise (Figure 4 and Table 3).

Microbial eukaryotic cell abundances were enriched within diffuse vent fluids (average of 230-620 cells ml^-1^); at minimum, there was an over two-fold higher concentration of eukaryotic cells in diffuse fluids compared to non-vent seawater (90-150 cell ml^-1^). This trend parallels observations of bacterial and archaeal abundances at hydrothermal vents [54] as well as patterns of protistan community diversity and species richness, confirming previous hypotheses regarding microeukaryotic vent populations [3, 6]. These findings demonstrate how active diffuse flow produces, attracts, and supports a greater biomass and diversity of deep-sea microorganisms. Outside the range of direct diffuse flow, plume and background samples had comparable eukaryotic cell counts to previous records from mesopelagic depths, which ranged from 74 to 400 cells ml^-1^ [31, 52, 55, 56]. Furthermore, this work contributes to growing evidence that deep-sea vents supply a substantial amount of labile carbon to the deep sea [15, 57].

Biomass estimates derived from two separate carbon conversion strategies revealed that deep-sea microeukaryotes make up a large amount of the carbon pool present at the diffuse vent fluid-seawater interface, which represents a significant energy resource for other vent organisms (Figure 4). In Pernice et al. [31], protistan biomass was found to decrease with depth, from 0.28 µg C L^-1^ at 200-450 m to 0.05 µg C L^-1^ at 1401-4000 m. These values are lower than what was found in the non-vent environment at Mid-Cayman Rise (2.5 - 12.9 µg C L^-1^; Table 1), which may be explained by the proximity of the hydrothermal vent to the plume and deep seawater in this study compared to the meso-to bathypelagic environment sampled in Pernice et al [31]. The amount of carbon represented by the hydrothermal vent protistan community is also significant as it demonstrates that protists can serve as a food resource to higher order consumers (Table 2) [15]. Studies of larger macrofauna at vent sites suggest that their diets include isotopically-varied food sources [58], which includes microbial eukaryotes [59].

Paired molecular analyses show that ciliates and flagellates make up a large proportion of the grazer community (Figure 3b), which is consistent with previous work [6, 10, 11]. The higher biomass measured within vent fluids may be explained by the increase in larger eukaryotes, such as ciliates (Table 2; Figure S2). While flagellates are documented as key grazers throughout the deep sea and mesopelagic [31], the increase in prey availability and resources at vent sites can sustain larger ciliate cells. This corroborates observed increases in biomass detected at vents compared to non-vent environments in this study, and in Pachiadaki et al. [52], where there was an increase in ciliates, relative to flagellates, at a biological hotspot in the deep sea (halocline).

Diffuse vent sites with the highest recorded temperature typically included the highest concentrations of microeukaryotic and microbial prey cells, and subsequent grazing rates (Figure 2). The mixing of heated, end-member hydrothermal fluid and cold oxygenated seawater generated an increase in available oxidants and reductants for increased microbial metabolic activity; this ultimately enhances chemosynthetic productivity at diffuse vent sites [60]. This is especially true at ultramafic sites, like Von Damm, where both subsurface abiotic and biotic carbon synthesis within mixing vent fluid contributes to a higher availability of labile carbon [15, 46, 60]. Thus, between Piccard and Von Damm, we originally expected the highest grazing rates to be at Von Damm. However, average grazing rates at Piccard were higher relative to Von Damm; we attributed this to a temperature limitation; the highest temperatures at Von Damm (peak of 121°C during collection) appeared to limit cell abundances, causing a plateau (Figure 2). Factors controlling protistan grazing pressure are often found to be temperature and abundances of predators and prey, similar to what we found, but temperature limited on grazing capacity have also been observed [61]. We emphasize the role that temperature plays in this study, as other environmental parameters did not demonstrate as clear of a trend (Figure S6); however, we acknowledge that co-varying parameters or other cryptic processes likely contribute to microbial food web dynamics in the deep sea (Figure 2).

### Broader Implications of microbially-mediated C-flux

The influence of hydrothermal vent food webs extends beyond the local vent region, providing resources to the surrounding deep-sea ecosystem that may outweigh export from the sea surface [59, 62, 63]. Our results demonstrate that the amount of carbon biomass stored within the prokaryotic and eukaryotic microbiota is substantial, and can vary as a factor of hydrothermal vent geochemistry, community composition, and temperature. Using carbon as the currency in Figure 4, we illustrate the potential range of carbon biomass and trophic transfer, beginning with chemosynthetic primary production, to consumption, and extending to unconstrained routes of carbon export.

The effect of increased microbial interactions and a chemosynthetically-sourced microbial food web within mixed diffuse fluid flow and deep seawater also reaches the rising hydrothermal vent plume. Within the hydrothermal plume, entrained diffuse fluid and deep seawater support its own biological habitat [64]. Within dispersing plumes, Bennett et al. [57] and Bennett et al. [15] detected elevated organic carbon production, above East Pacific Rise (EPR) and Mid-Cayman Rise vent fields, respectively. While increased particulate organic carbon (POC) and dissolved organic carbon (DOC) was observed at EPR, the POC was disproportionately higher relative to DOC and to the background environment, leading the authors to hypothesize that microbially-mediated processes are responsible for carbon production [57]. Our findings strengthen this conclusion, where a productive microbial food web contributes to the repackaging of DOC into POC, ultimately contributing to increased carbon in the plume and export into the surrounding ocean.

### Summary

Our findings contribute novel estimates of deep-sea hydrothermal vent protistan populations and biomass; when combined with quantitative measurements of predation pressure we present a more comprehensive view of the deep-sea hydrothermal vent microbial food web. We demonstrate that best practice is to conduct experiments under *in situ* conditions. Considering the required access to technology and necessary time and effort to conduct *in situ* incubations in the deep sea, we also show that results from experiments at ambient pressure contribute meaningful observations. A larger implication of this work is that the hydrothermal vent-associated microbial food web plays a significant role in the broader deep-sea carbon budget, as export from hydrothermal vents can contribute a substantial amount of local carbon and other nutrient resources (Figure 4).

## Acknowledgements

Authors would like to acknowledge everyone who contributed to cruise operations and sample collection. We would like to thank the captain and crew of the RV Atlantis and the pilots and engineers for ROV Jason. Sample collection and processing would not have been possible without the support of Malayika Vincent, Jessica Frankle, and Eoghan Reeves. Funding support for the Mid-Cayman Rise expedition was provided by NSF OCE-1801036 (to S.Q.L), 1801205 to (J.S.S), NSF OCE-1816652, and NASA Planetary Science and Technology Through Analog Research (PSTAR) Program 80NSSC17K0252 (S.Q.L.). G.R.G was supported by NSF OCE-185100. V.P.E. was supported by NSF OCE-1737173. The NSF Center for Dark Energy Biosphere Investigations (C-DEBI) supported J.A.H. as well as S.K.H. through a C-DEBI Postdoctoral Fellowship (OCE-0939564). Support for sample processing and data analysis was also provided by the Charles E. Hollister Endowed Fund for Support of Innovative Research at Woods Hole Oceanographic Institution (WHOI). Research and analysis were also supported through NSF grants OCE-1947776 awarded to J.A.H., M.G.P., and V.P.E. and OCE-2327203 awarded to R.A.A., J.A.H., and S.K.H. CDEBI contribution ### (*tbd*).

## Competing Interests

Authors declare no competing interests.

## Data Availability Statement

All necessary data products and code to reproduce results can be found at https://shu251.github.io/midcayman-rise-microeuk/. Raw sequence data are available through NCBI, SRA BioProject accession number PRJNA802868.

---

**Figure S1.**
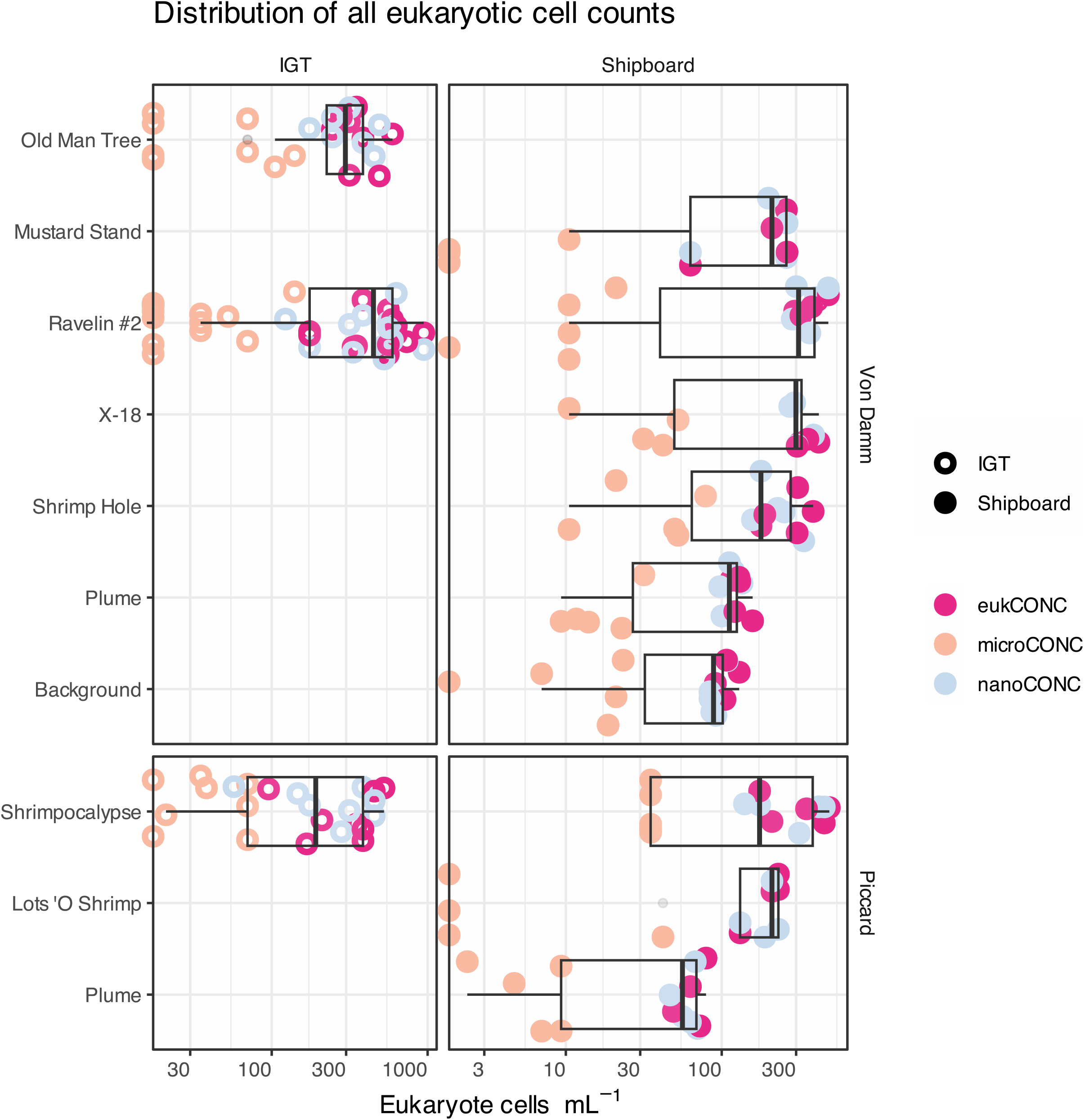
Eukaryotic cell counts from all grazing samples (x-axis is cells ml^-1^; log scale). Including separation of micro-and nano-size classes by color. Left panels represent cell counts from IGT experiments and right hand panels originate from shipboard samples.

**Figure S2.**
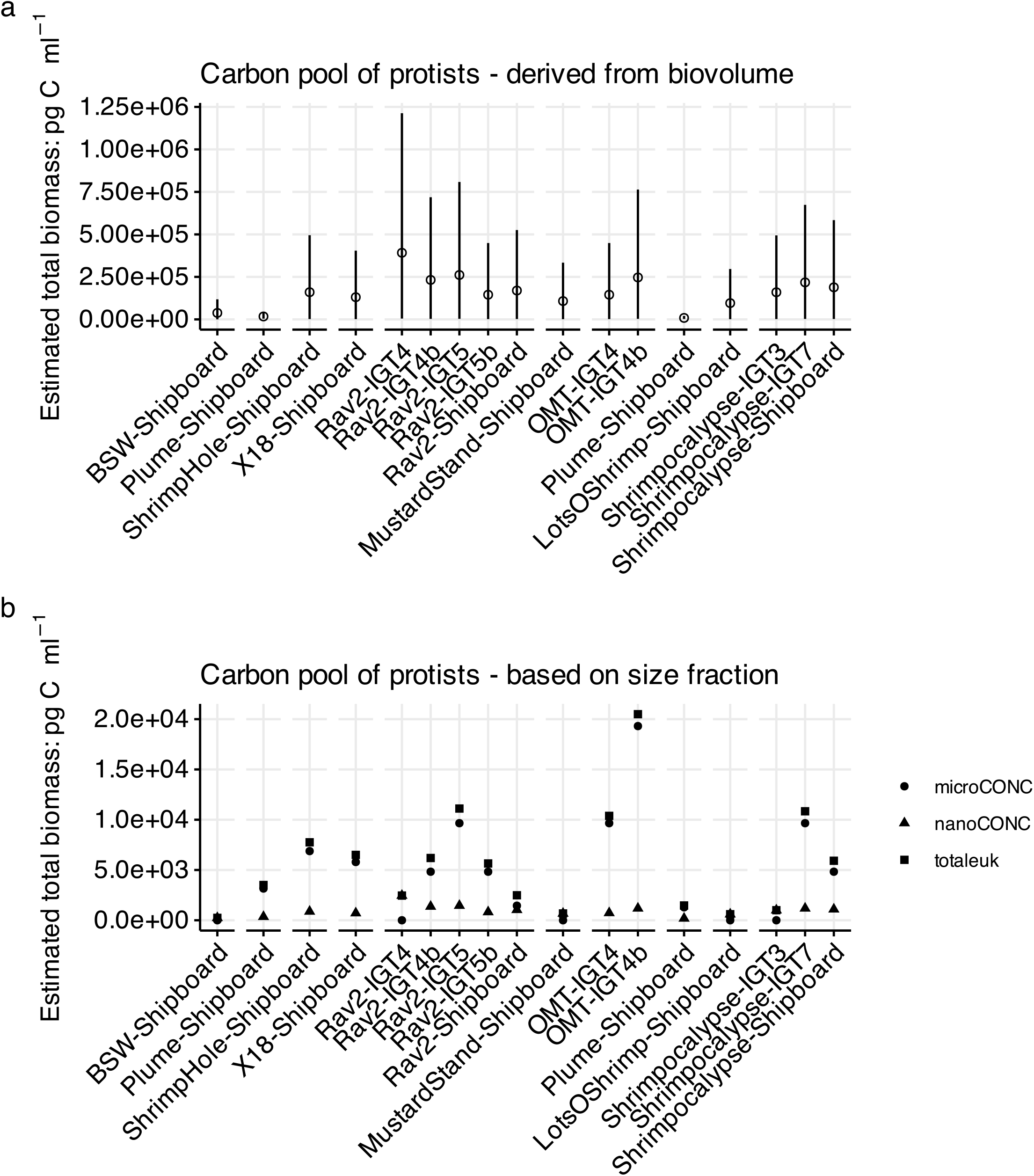
Estimated range and mean of protistan biomass use either (a) the biovolume of imaged cells together with a carbon conversion factor that incorporates a diverse group of protists, excluding diatoms [33] or (b) assigning the nano and micro size classes individual carbon conversion factors [34].

**Figure S3.**
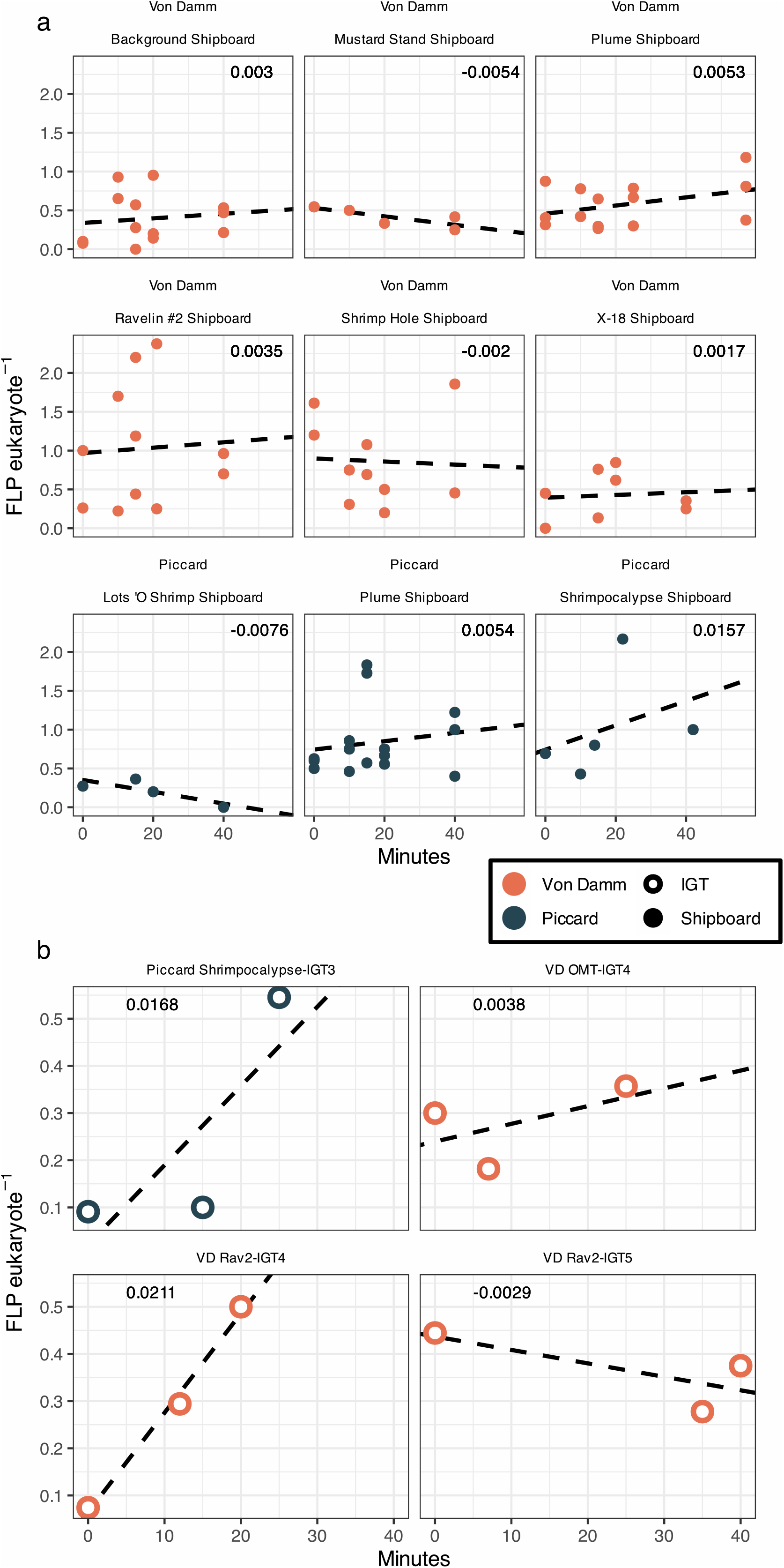
Results from each conducted grazing experiment, where experiment time is along the x-axis and the number of FLP eukaryotic cell^-1^ detected is noted at the y-axis. The number indicated at the top right hand corner of each panel indicates the calculated slope of the line. For values that are negative, this is noted as undetected and reflected as zero in the main results. For IGT samples, the final time point (T40) was removed, as the small incubation chamber size likely caused a bottle effect (Figure S4).

**Figure S4.**
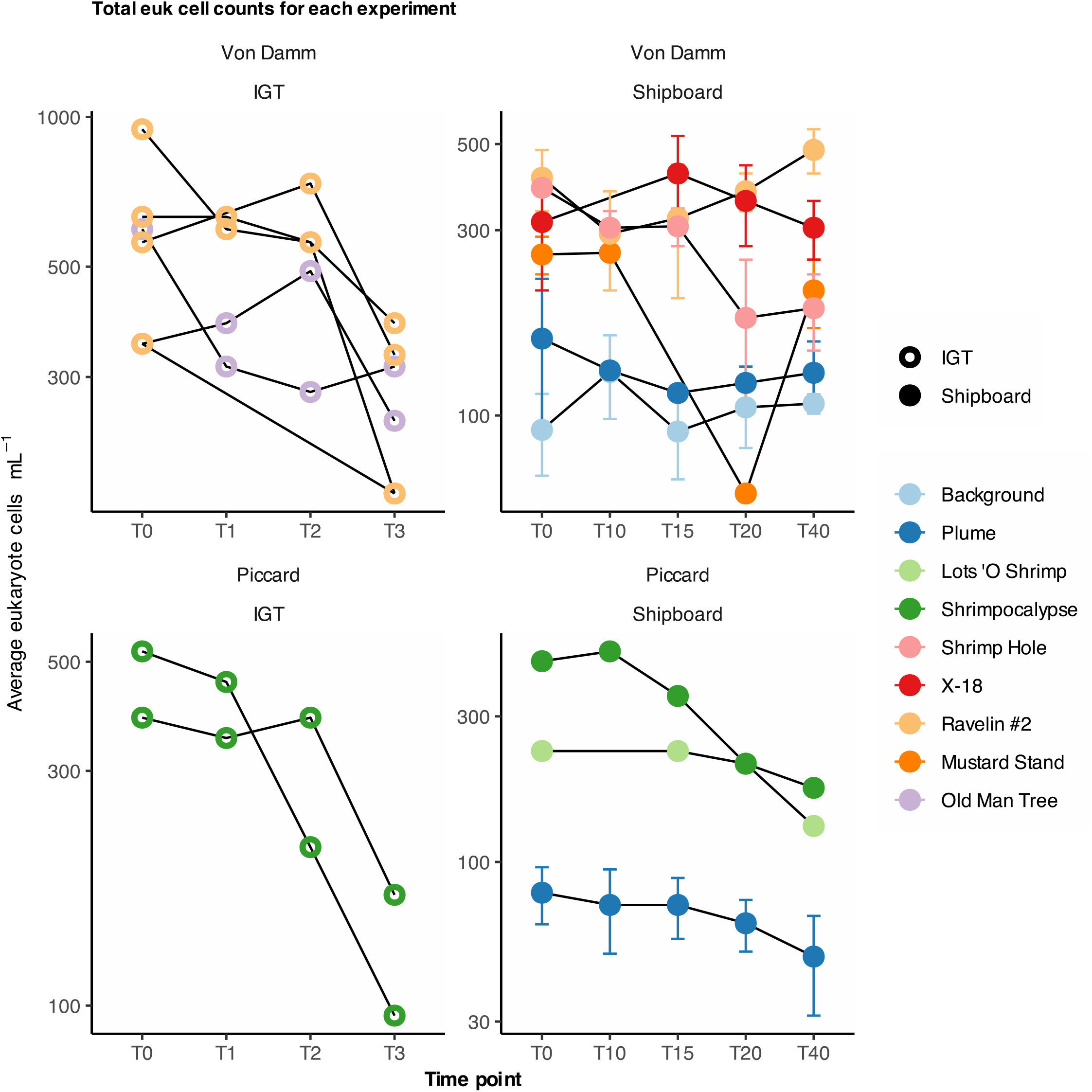
Total eukaryotic cells ml^-1^ at each time point for IGT (left) and shipboard (right) experiments. Note that the IGT experiments appear to have a bottle effect (smaller volume), so the final time point (T3 or approx. 40 minutes) was removed for downstream analyses (Figure S4).

**Figure S5.**
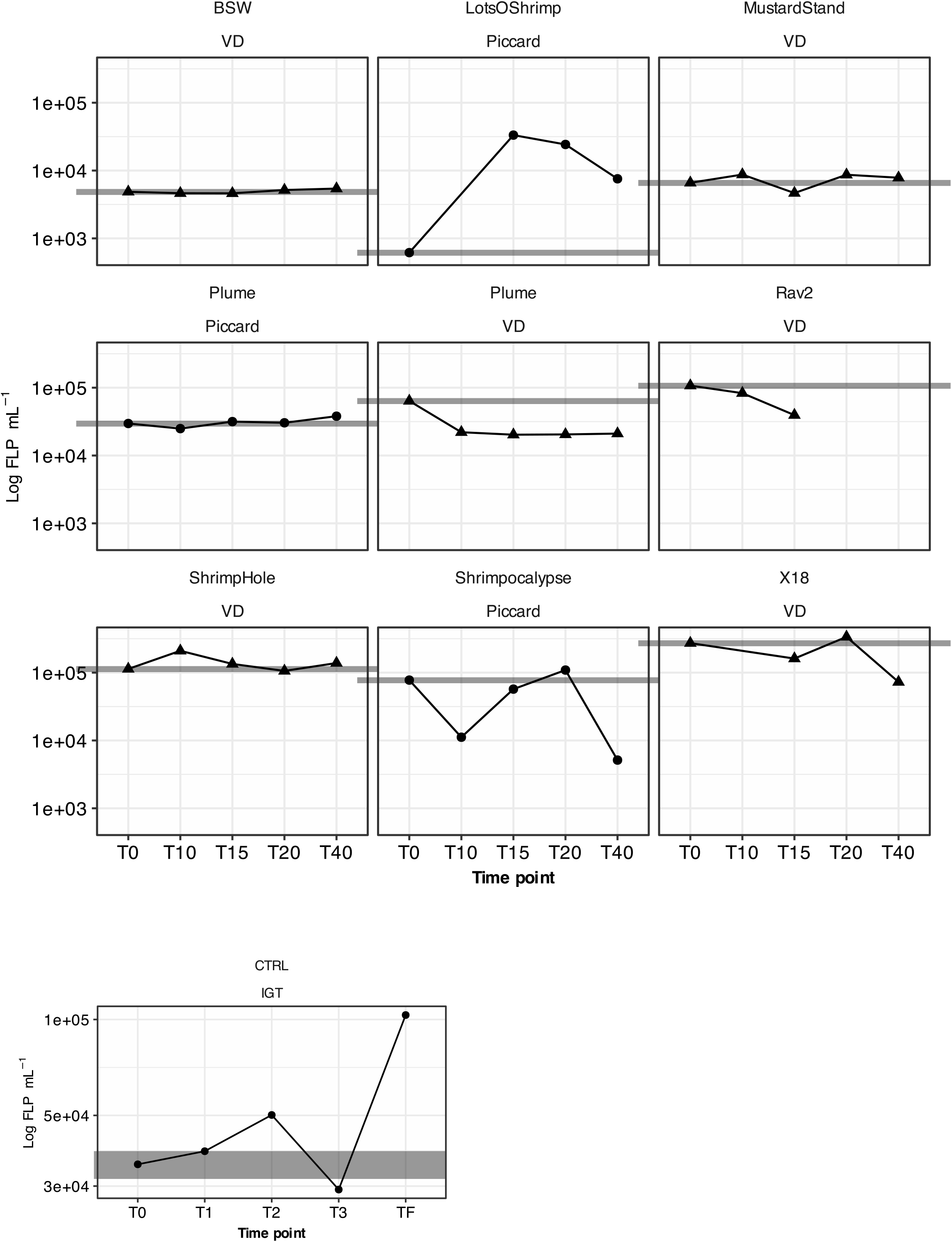
FLP control experiments. Ideally, FLP concentration in control experiments remains stable over time. For most experiments shown here, FLP concentration varied little or could be explained by insufficient mixing at time zero; this is shown when T0 is the outlier. For the IGT experiments, plume and background seawater was used to mimic the control experiment. Due to the nature of IGT sampling, control samples could not be run at the same time as the experimental.

**Figure S6.**
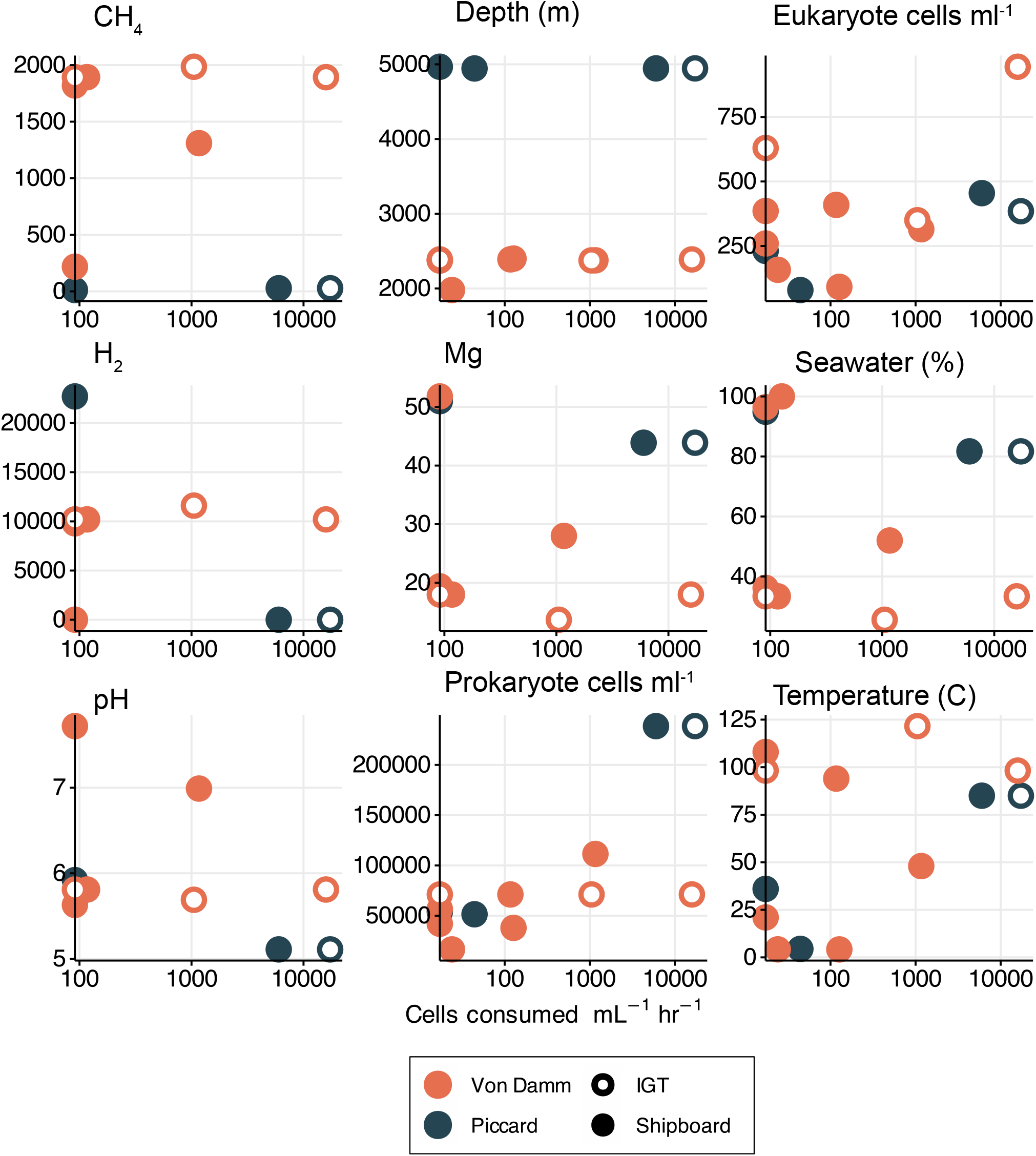
All environmental parameters (y-axes) vs. protistan grazing rate (cells consumed ml^-1^ hr^-1^ (x-axis). Each panel represents a separate environmental parameter (see Table S1 for units and values). Eukaryotic and prokaryotic cells ml^-1^ and temperature (°C) appeared to have a relationship with grazing rate and are investigated further in the main text.

**Figure S7.**
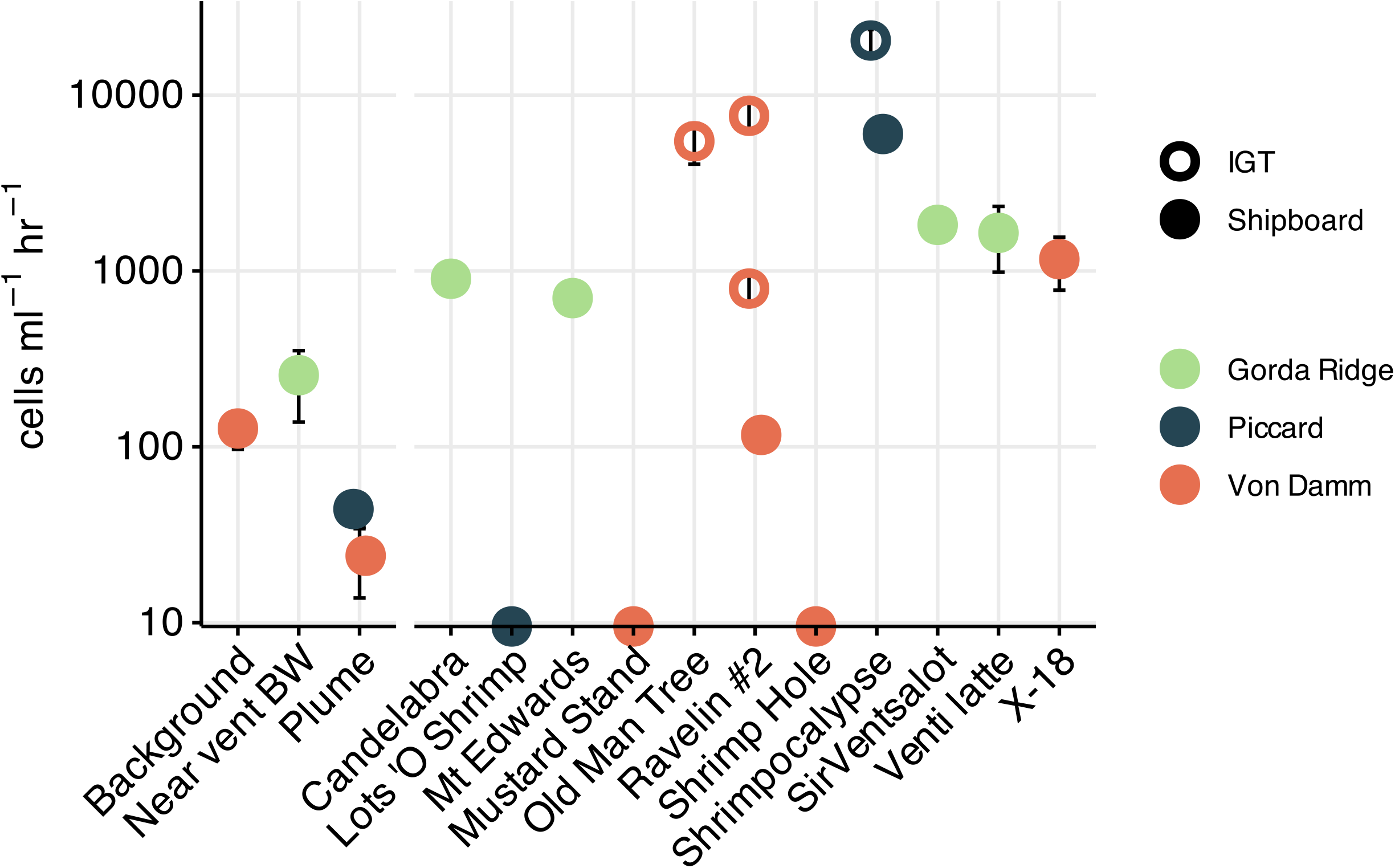
Comparison of grazing rates across three hydrothermal vent fields: Gorda Ridge [10], Von Damm, and Piccard (this study). Rates are expressed in log scale as the number of cells consumed by protistan predators ml^-1^ hr^-1^. Symbol color denotes vent field, filled in circles are derived from shipboard experiments or samples, while circle outlines represent results from IGT experiments.

**Figure S8.**
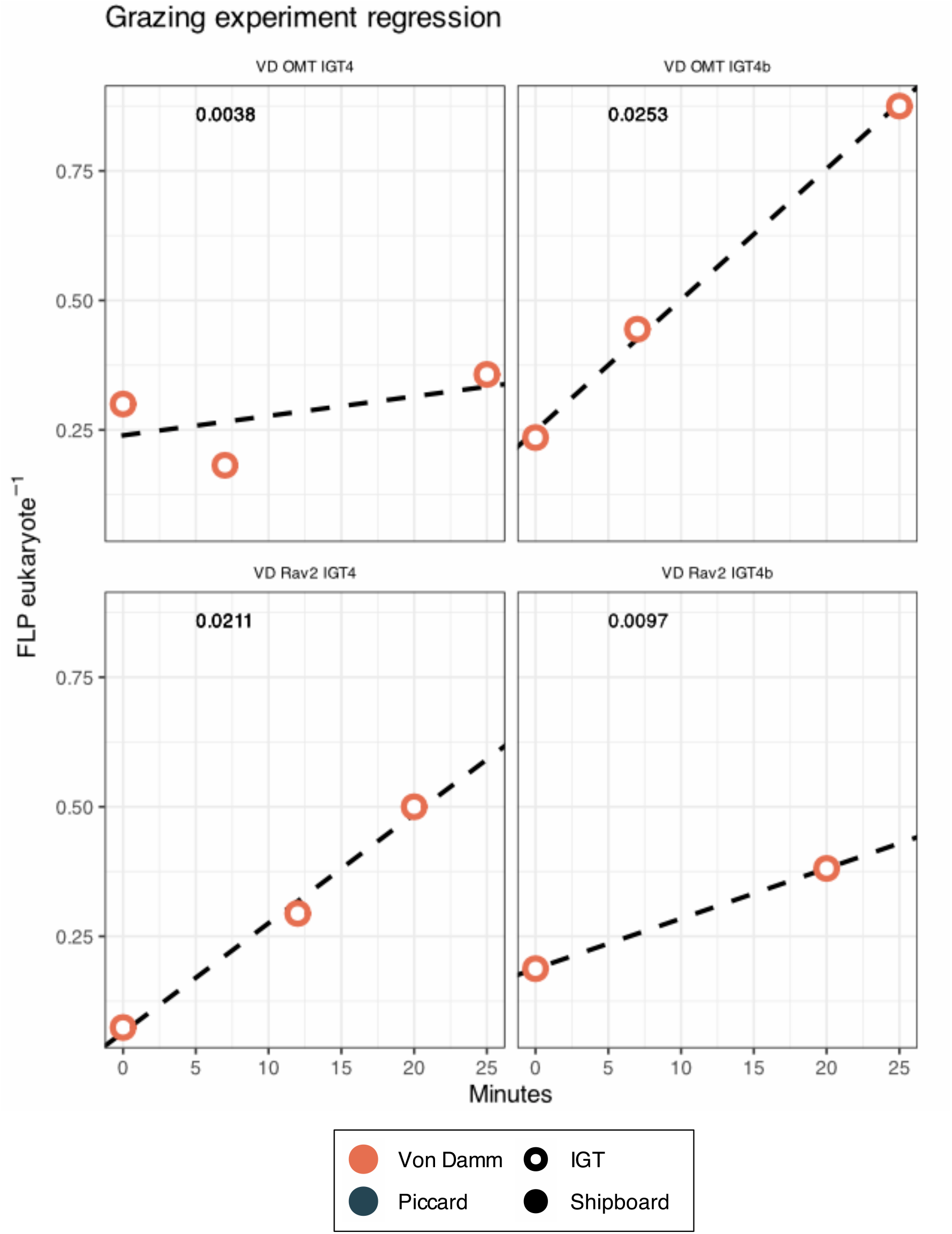
Technical replicates for Old Man Tree (top panel) and Ravelin #2 (bottom panels). Since biological replicates were difficult to obtain for the IGT experiments, two halves of a filter were counted separately as technical replicates. On the right side are the results in this study and the left side reports results from “IGTxb” technical replicates. Together with grazing rates that were in range of another study (Figure S6), we can provide additional support in our findings.

**Figure S9.**
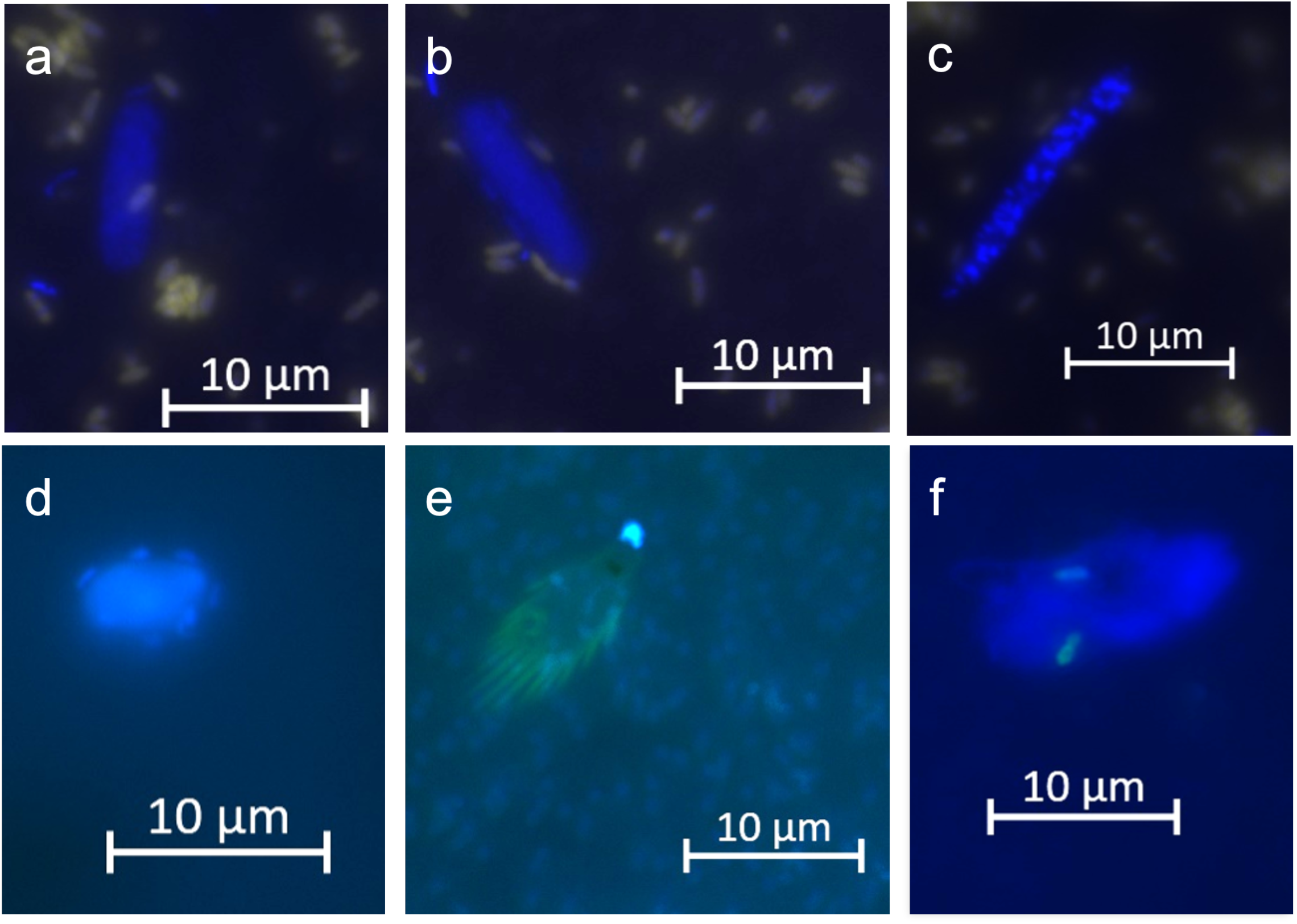
Microeukaryotic cells captured with epifluorescence (DAPI filter) from FLP uptake experiments conducted in this study. All images are derived from shipboard experiments and scale bars are located within each panel. **(a-c)** Eukaryotic cells from grazing experiments conducted at vent site X-18. The cell in panel (a) contained a consumed FLP cell. Imaged cells from **(d)** Mustard Stand, **(e)** Shrimpocalypse, and **(f)** the plume at Von Damm. Two FLP cells are imaged inside the eukaryotic cell in (f).

**Table S1.** Metadata for all samples depth (meters), maximum temperature (°C) at time of collection, pH, estimated percent seawater in diffuse fluids (%), pH, magnesium (Mg mmol/L or mM), dissolved hydrogen (H2 μmol/L or μM), total dissolved hydrogen sulphide (H2S mmol/L or mM), and methane (CH4 μmol/L or μM), concentration of microorganisms (cells/ml), location, and fluid origin type.

**Table S2.** Complete experiment details for all shipboard and IGT grazing assays, including ROV Jason number, time points, number of replicates, controls, and volumes. Table also reports major results from cell counts, grazing rate, clearance rate, and percent bacterial turnover percentage.

**Table S3.** List of all variables and calculations for this study, related to grazing experiments (top) or estimates of carbon biomass. In addition to the variable used in associated R code, the equations, units, and references are also listed.

**Table S4.** Calculation of microbial eukaryotic biovolume from Hillebrand et al. [32] and estimate of carbon content cell^-1^ based on Menden-Deuer & Lessard 2000 [33].

**Table S5.** Estimates of protistan biomass in pg C cell^-1^ derived from either biovolume or size fractionation approach. In addition to total eukaryotic biomass, table lists estimates based on size fraction, nano-(<20µm) versus micro-(≥20µm).

## Notes

https://shu251.github.io/midcayman-rise-microeuk/

### Competing Interest Statement

The authors have declared no competing interest.

## References

1. Van Dover CL. The Ecology of Deep-Sea Hydrothermal Vents. 2021. Princeton University Press.

2. Tunnicliffe V, McArthur AG, McHugh D. A Biogeographical Perspective of the Deep-Sea Hydrothermal Vent Fauna. In: Blaxter JHS, Southward AJ, Tyler PA (eds). Advances in Marine Biology. 1998. Academic Press, pp 353–442.

3. Murdock SA, Juniper SK. Hydrothermal vent protistan distribution along the Mariana arc suggests vent endemics may be rare and novel. Environ Microbiol 2019; 21: 3796–3815.

4. Edgcomb VP, Kysela DT, Teske A, Gomez A de V, Sogin ML. Benthic Eukaryotic Diversity in the Guaymas Basin Hydrothermal Vent Environment. Proc Natl Acad Sci U S A 2002; 99: 7658–7662.

5. Lopez-Garcia P, Philippe H, Gail F, Moreira D. Autochthonous eukaryotic diversity in hydrothermal sediment and experimental microcolonizers at the Mid-Atlantic Ridge. Proceedings of the National Academy of Sciences 2003; 100: 697–702.

6. Hu SK, Smith AR, Anderson RE, Sylva SP, Setzer M, Steadmon M, et al. Globally-distributed microbial eukaryotes exhibit endemism at deep-sea hydrothermal vents. Mol Ecol 2022.

7. Huber JA, Butterfield DA, Baross JA. Diversity and distribution of subseafloor Thermococcales populations in diffuse hydrothermal vents at an active deep-sea volcano in the northeast Pacific Ocean: Diversity of Subseafloor Thermococcales. J Geophys Res Biogeosci 2006; 111.

8. Akerman NH, Butterfield DA, Huber JA. Phylogenetic diversity and functional gene patterns of sulfur-oxidizing subseafloor Epsilonproteobacteria in diffuse hydrothermal vent fluids. Front Microbiol 2013; 4.

9. Thomas E, Anderson RE, Li V, Rogan LJ, Huber JA. Diverse Viruses in Deep-Sea Hydrothermal Vent Fluids Have Restricted Dispersal across Ocean Basins. mSystems 2021; 6: e0006821.

10. Hu SK, Herrera EL, Smith AR, Pachiadaki MG, Edgcomb VP, Sylva SP, et al. Protistan grazing impacts microbial communities and carbon cycling at deep-sea hydrothermal vents. Proc Natl Acad Sci U S A 2021; 118.

11. Pasulka A, Hu SK, Countway PD, Coyne KJ, Cary SC, Heidelberg KB, et al. SSU rRNA Gene Sequencing Survey of Benthic Microbial Eukaryotes from Guaymas Basin Hydrothermal Vent. J Eukaryot Microbiol 2019; jeu.12711.

12. Moreira D. Are hydrothermal vents oases for parasitic protists? Trends Parasitol 2003; 19: 556–558.

13. Rotterová J, Edgcomb VP, Čepička I, Beinart R. Anaerobic ciliates as a model group for studying symbioses in oxygen-depleted environments. J Eukaryot Microbiol 2022; e12912.

14. Beinart RA, Beaudoin DJ, Bernhard JM, Edgcomb VP. Insights into the metabolic functioning of a multipartner ciliate symbiosis from oxygen-depleted sediments. Mol Ecol 2018; 27: 1794–1807.

15. Bennett SA, Coleman M, Huber JA, Reddington E, Kinsey JC, McIntyre C, et al. Trophic regions of a hydrothermal plume dispersing away from an ultramafic-hosted vent-system: Von Damm vent-site, Mid-Cayman Rise. Geochem Geophys Geosyst 2013; 14: 317–327.

16. Van Dover CL, Fry B. Microorganisms as food resources at deep-sea hydrothermal vents. Limnol Oceanogr 1994; 39: 51–57.

17. Edgcomb VP, Taylor C, Pachiadaki MG, Honjo S, Engstrom I, Yakimov M. Comparison of Niskin vs. in situ approaches for analysis of gene expression in deep Mediterranean Sea water samples. Deep Sea Res Part 2 Top Stud Oceanogr 2016; 129: 213–222.

18. Sievert SM, Vetriani C. Chemoautotrophy at deep-sea vents: past, present, and future. Oceanography 2012; 25: 218–233.

19. Edgcomb V, Orsi W, Taylor GT, Vdancy P, Taylor C, Suarez P, et al. Accessing marine protists from the anoxic Cariaco Basin. ISME J 2011; 5: 1237–1241.

20. Breier JA, Sheik CS, Gomez-Ibanez D, Sayre-McCord RT, Sanger R, Rauch C, et al. A large volume particulate and water multi-sampler with in situ preservation for microbial and biogeochemical studies. Deep Sea Res Part I 2014; 94: 195–206.

21. Fortunato CS, Butterfield DA, Larson B, Lawrence-Slavas N, Algar CK, Zeigler Allen L, et al. Seafloor Incubation Experiment with Deep-Sea Hydrothermal Vent Fluid Reveals Effect of Pressure and Lag Time on Autotrophic Microbial Communities. Appl Environ Microbiol 2021; 87.

22. Seewald JS, Doherty KW, Hammar TR, Liberatore SP. A new gas-tight isobaric sampler for hydrothermal fluids. Deep Sea Res Part I 2002; 49: 189–196.

23. McNichol J, Sylva SP, Thomas F, Taylor CD, Sievert SM, Seewald JS. Assessing microbial processes in deep-sea hydrothermal systems by incubation at in situ temperature and pressure. Deep Sea Res Part I 2016; 115: 221–232.

24. Lang SQ, Benitez-Nelson B. Hydrothermal Organic Geochemistry (HOG) sampler for deployment on deep-sea submersibles. Deep Sea Res Part I 2021; 173: 103529.

25. Sherr EB, Sherr BF. Protistan Grazing Rates via Uptake of Fluorescently Labeled Prey. Handbook of methods in aquatic microbial 1993.

26. Caron DA. Protistan herbivory and bacterivory. Methods in Microbiology. 2001., 289–315

27. Sherr BF, Sherr EB, Fallon RD. Use of monodispersed, fluorescently labeled bacteria to estimate in situ protozoan bacterivory. Appl Environ Microbiol 1987; 53: 958–965.

28. Trembath-Reichert E, Butterfield DA, Huber JA. Active subseafloor microbial communities from Mariana back-arc venting fluids share metabolic strategies across different thermal niches and taxa. ISME J 2019; 13: 2264–2279.

29. Unrein F, Massana R, Alonso-Sáez L, Gasol JM. Significant year-round effect of small mixotrophic flagellates on bacterioplankton in an oligotrophic coastal system. Limnol Oceanogr 2007; 52: 456–469.

30. Schneider CA, Rasband WS, Eliceiri KW. NIH Image to ImageJ: 25 years of image analysis. Nat Methods 2012; 9: 671–675.

31. Pernice MC, Forn I, Gomes A, Lara E, Alonso-Sáez L, Arrieta JM, et al. Global abundance of planktonic heterotrophic protists in the deep ocean. ISME J 2015; 9: 782–792.

32. Hillebrand H, Dürselen C-D, Kirschtel D, Pollingher U, Zohary T. Biovolume calculation for pelagic and benthic microalgae. J Phycol 1999; 35: 403–424.

33. Menden-Deuer S, Lessard EJ. Carbon to volume relationships for dinoflagellates, diatoms, and other protist plankton. Limnol Oceanogr 2000; 45: 569–579.

34. Caron DA, Dam HG, Kremer P, Lessard EJ, Madin LP, Malone TC, et al. The contribution of microorganisms to particulate carbon and nitrogen in surface waters of the Sargasso Sea near Bermuda. Deep Sea Res Part I 1995; 42: 943–972.

35. Choi JW, Stoecker DK. Effects of fixation on cell volume of marine planktonic protozoa. Appl Environ Microbiol 1989; 55: 1761–1765.

36. Verity PG, Robertson CY, Tronzo CR, Andrews MG, Nelson JR, Sieracki ME. Relationships between cell volume and the carbon and nitrogen content of marine photosynthetic nanoplankton. Limnol Oceanogr 1992; 37: 1434–1446.

37. Morono Y, Terada T, Nishizawa M, Ito M, Hillion F, Takahata N, et al. Carbon and nitrogen assimilation in deep subseafloor microbial cells. Proc Natl Acad Sci U S A 2011; 108: 18295–18300.

38. Loferer-Krößbacher M, Klima J, Psenner R. Determination of Bacterial Cell Dry Mass by Transmission Electron Microscopy and Densitometric Image Analysis. Appl Environ Microbiol 1998; 64: 688–694.

39. Stoeck T, Zuendorf A, Breiner H-W, Behnke A. A Molecular Approach to Identify Active Microbes in Environmental Eukaryote Clone Libraries. Microb Ecol 2007; 53: 328–339.

40. Bolyen E, Rideout JR, Dillon MR, Bokulich NA, Abnet CC, Al-Ghalith GA, et al. Reproducible, interactive, scalable and extensible microbiome data science using QIIME 2. Nat Biotechnol 2019; 37: 852–857.

41. Guillou L, Bachar D, Audic S, Bass D, Berney C, Bittner L, et al. The Protist Ribosomal Reference database (PR2): a catalog of unicellular eukaryote Small Sub-Unit rRNA sequences with curated taxonomy. Nucleic Acids Res 2012; 41: D597–D604.

42. Vaulot D. pr2database/pr2database: PR2 version 4.14.0. 2021.

43. Martin BD, Witten D, Willis AD. Corncob: count regression for correlated observations with the beta-binomial. corncob: Count Regression for Correlated Observations with the Beta-binomial R 2020.

44. Connelly DP, Copley JT, Murton BJ, Stansfield K, Tyler PA, German CR, et al. Hydrothermal vent fields and chemosynthetic biota on the world’s deepest seafloor spreading centre. Nat Commun 2012; 3: 620.

45. Kinsey JC, German CR. Sustained volcanically-hosted venting at ultraslow ridges: Piccard Hydrothermal Field, Mid-Cayman Rise. Earth Planet Sci Lett 2013; 380: 162–168.

46. McDermott JM, Seewald J, Reeves EP, German CR, Sylva SP, Klein F. Abundance of volatile and organic species in intermediate temperature fluids from the Von Damm and Piccard deep sea hydrothermal fields, Mid-Cayman Rise. 2012. p OS22B–07.

47. McDermott JM, Sylva SP, Ono S, German CR, Seewald JS. Geochemistry of fluids from Earth’s deepest ridge-crest hot-springs: Piccard hydrothermal field, Mid-Cayman Rise. Geochim Cosmochim Acta 2018; 228: 95–118.

48. Gong J, Dong J, Liu X, Massana R. Extremely High Copy Numbers and Polymorphisms of the rDNA Operon Estimated from Single Cell Analysis of Oligotrich and Peritrich Ciliates. Protist 2013; 164: 369–379.

49. Gloor GB, Macklaim JM, Pawlowsky-Glahn V, Egozcue JJ. Microbiome Datasets Are Compositional: And This Is Not Optional. Front Microbiol 2017; 8.

50. Turley CM, Carstens M. Pressure tolerance of oceanic flagellates: implications for remineralization of organic matter. Deep Sea Res A 1991; 38: 403–413.

51. Medina LE, Taylor CD, Pachiadaki MG, Henríquez-Castillo C, Ulloa O, Edgcomb VP. A Review of Protist Grazing Below the Photic Zone Emphasizing Studies of Oxygen-Depleted Water Columns and Recent Applications of In situ Approaches. Frontiers in Marine Science 2017; 4.

52. Pachiadaki MG, Taylor C, Oikonomou A, Yakimov MM, Stoeck T, Edgcomb V. In situ grazing experiments apply new technology to gain insights into deep-sea microbial food webs. Deep Sea Res Part 2 Top Stud Oceanogr 2016; 129: 223–231.

53. Connell PE, Campbell V, Gellene AG, Hu SK, Caron DA. Planktonic food web structure at a coastal time-series site: II. Spatiotemporal variability of microbial trophic activities. Deep Sea Res Part I 2017; 121: 210–223.

54. Fortunato CS, Larson B, Butterfield DA, Huber JA. Spatially distinct, temporally stable microbial populations mediate biogeochemical cycling at and below the seafloor in hydrothermal vent fluids: Microbial genomics at axial seamount. Environ Microbiol 2018; 20: 769–784.

55. Rocke E, Pachiadaki MG, Cobban A, Kujawinski EB, Edgcomb VP. Protist Community Grazing on Prokaryotic Prey in Deep Ocean Water Masses. PLoS One 2015; 10: e0124505.

56. Cho B, Na S, Choi D. Active ingestion of fluorescently labeled bacteria by mesopelagic heterotrophic nanoflagellates in the East Sea, Korea. Mar Ecol Prog Ser 2000; 206: 23–32.

57. Bennett SA, Statham PJ, Green DRH, Le Bris N, McDermott JM, Prado F, et al. Dissolved and particulate organic carbon in hydrothermal plumes from the East Pacific Rise, 9 degree 50’N. Deep Sea 2011.

58. Bergquist D, Eckner J, Urcuyo I, Cordes E, Hourdez S, Macko S, et al. Using stable isotopes and quantitative community characteristics to determine a local hydrothermal vent food web. Mar Ecol Prog Ser 2007; 330: 49–65.

59. Govenar B. Energy transfer through food webs at hydrothermal vents: Linking the lithosphere to the biosphere. Oceanography 2012; 25: 246–255.

60. Reveillaud J, Reddington E, McDermott J, Algar C, Meyer JL, Sylva S, et al. Subseafloor microbial communities in hydrogen-rich vent fluids from hydrothermal systems along the Mid-Cayman Rise: Subseafloor microbes at Mid-Cayman Rise. Environ Microbiol 2016; 18: 1970–1987.

61. Vaqué D, Gasol JM, Marrasé C. Grazing rates on bacteria: the significance of methodology and ecological factors. Mar Ecol Prog Ser 1994; 109: 263–274.

62. Levin LA, Baco AR, Bowden DA, Colaco A, Cordes EE, Cunha MR, et al. Hydrothermal Vents and Methane Seeps: Rethinking the Sphere of Influence. Front Mar Sci 2016; 3.

63. Bell JB, Woulds C, van Oevelen D. Hydrothermal activity, functional diversity and chemoautotrophy are major drivers of seafloor carbon cycling. Sci Rep 2017; 7: 12025.

64. Dick GJ, Anantharaman K, Baker BJ, Li M, Reed DC, Sheik CS. The microbiology of deep-sea hydrothermal vent plumes: ecological and biogeographic linkages to seafloor and water column habitats. Front Microbiol 2013; 4: 124.

