## supplementary information for "Microbial eukaryotic predation pressure and biomass at deep-sea hydrothermal vents: Implications for deep-sea carbon cycling"

### I. IGT Technical replication

Only 1 IGT experiment was permitted per ROV Dive, thus biological replicates for IGTs were not carried out. Replication across separate dives or to compare with shipboard experiments was attempted for Ravelin #2, Old Man Tree, and Shrimpocalypse (Table 1, Table S1), but some of the grazing experiments did not work or were uncountable due to particulates, thus compromising an accurate count. For replication, biological replicates between IGT and shipboard were done for Ravelin #2 and Shrimpocalypse (reported in main text, see Table 1, IGT7). To further ground truth our methods, we also re-counted samples as technical replicates from the IGT experiments (denoted with "b" in Figure S7).

We saw consistent patterns across IGT and shipboard biological replicates, and technical replicates were comparable. However, due to bottle effects, we did not want to average across biological and technical replicates, thus these are kept separate in our analysis.

### II. Eukaryotic cell size and biomass

Due to the cell shrinkage from fixation and depressurization from the deep sea, we determined that microscopic designations between micro (> 20μm) and nano (<20 μm) were not accurate (Choi and Stoecker 1989; Edgcomb et al. 2016). While results from micro- and nano-eukaryote counts are reported as supplemental, major interpretations from the study are based on total eukaryote concentrations and biomass, which is the sum of the micro and nano size fractions.

Examples of eukaryotic cells captured with epifluorescence can be found in Figure S8.
